# Despotism promotes cooperation through enhanced interdependencies in non-human primate societies

**DOI:** 10.1101/2025.03.27.645840

**Authors:** Debottam Bhattacharjee, Tonko W. Zijlstra, Tom S. Roth, Elena Belli, Sophie Calis, Paula Escriche Chova, Eythan Cousin, Jolanda A. de Jong, Edwin J.A.M. de Laat, Aníta Rut Guðjónsdóttir, Karline R.L. Janmaat, Elja J. Jeunink, Charlotte E. Kluiver, Penny E.N. Kuijer, Esmee Middelburg, Lena S. Pflüger, Veera I. Schroderus, Eva S.J. van Dijk, Jonas Verspeek, Sophie Waasdorp, Adam N. Zeeman, Elisabeth H.M. Sterck, Edwin J.C. van Leeuwen, Jorg J.M. Massen

## Abstract

The evolutionary mechanisms of cooperation are well-studied, yet what motivates individuals to cooperate remains unclear. Although conventionally associated with enhanced social tolerance (*self-domestication hypothesis*) and prosociality (*cooperative-breeding hypothesis*), individuals belonging to species that are neither self-domesticated nor cooperatively-breeding also cooperate. An overarching *interdependency hypothesis* posits that inter-individual dependencies promote cooperation, but comparative evidence is lacking. We experimentally studied cooperation, prosociality, and tolerance in six macaque species along a ‘despotic-egalitarian’ gradient. Within-group cooperation was higher in despotic than egalitarian societies yet restricted to few partners. Prosociality, kinship, and tolerance positively predicted cooperation success. Agent-based models consistently showed that despotic societies have fewer but more stable social bonds and, thus, higher interdependencies than egalitarian ones. Our results suggest that interdependencies facilitate the emergence and maintenance of cooperation.

## Introduction

Cooperation – collaborative efforts of two or more individuals to achieve a common goal – helps maintain societies and provides benefits to group members (*1*, *2*). Existing theoretical and empirical frameworks provide evolutionary explanations of kin- (through kin selection) and non-kin- (through reciprocal altruism) cooperation in animals (*3*, *4*). Yet, what motivates individuals to cooperate and how they choose reliable partners during cooperation remains unclear at the proximate level. Such motivations are fundamental to the emergence and sustenance of cooperation (*5*). Conventionally, cooperation has been linked to societies that exhibit enhanced social tolerance (*self-domestication hypothesis,* (*6*)) and prosociality (*cooperative breeding hypothesis,* (*7*)). However, growing empirical research shows that individuals in several species that are neither self-domesticated nor cooperatively breeding exhibit prosocial and cooperative tendencies, too (*8*–*15*). Building on the idea that interdependence – an individual’s stake in another – has a considerable role in the evolution of human cooperation (*16*, *17*), an overarching *interdependency hypothesis* may explain what might cause these species, and others in general, to show cooperative tendencies (*13*). This hypothesis suggests that interdependencies at the group level, for example, strength in numbers in colonially nesting species or allomaternal care in cooperatively breeding species (19), can lead to enhanced within-group social tolerance and promote indiscriminate prosociality and cooperation. It further emphasizes that at the dyadic level, preferential strong associations, nepotistic biases, and reliance on coalitions may result in selectively enhanced tolerance, promoting discriminate prosociality and cooperation, particularly in less tolerant or despotic societies (*12*, *13*). Studying the role of interdependencies in cooperation requires a comparative approach that includes closely related species with varying social tolerance, nepotistic, and coalitionary tendencies, but comparative empirical research remains scarce despite the potential to yield crucial insights into the evolution of cooperation.

Among group-living non-human primates, the genus *Macaca* consists of ∼22 species that have similar social organizations and group compositions but strikingly different levels of social tolerance, dominance hierarchy steepness, and nepotistic biases (*18*). This provides an excellent scope for comparative research to investigate the role of interdependencies in promoting cooperation. Macaque societies, primarily focusing on adult females, show a suite of these (often) co-varying traits and are traditionally categorized into four grades of ‘social styles’ (also known as tolerance grades) along a *despotic-egalitarian gradient* (*19*). Typically, despotic societies (e.g., Grades 1 and 2) are considered to show steeper hierarchies, more frequent aggressive interactions, and lower group-level tolerance than egalitarian societies (e.g., Grades 3 and 4) (*18*, *20*–*23*). Therefore, if group-level tolerance is a driver of cooperation, egalitarian macaque societies are expected to show higher (indiscriminate) cooperative tendencies than despotic macaque societies. By contrast, despotic societies show strong kin bonds and nepotistic biases, especially among females of the same matriline (*18*, *19*, *21*, *24*). Non-kin strong bonds may also form in despotic societies, primarily through friendships and coalitions (*12*, *25*, *26*), benefiting individuals during (post-)conflicts and helping to increase their social ranks. Such advantageous yet selective preferences among both kin- and non-kin group members can indicate the presence of strong interdependencies, leading to enhanced dyadic tolerance and (discriminate) prosocial- and cooperative-tendencies (*13*).

We studied 13 groups of macaques belonging to six species representing all four tolerance grades to test the predictions of the interdependency hypothesis (**Fig. 1A**). We conducted >29,800 minutes of observations and implemented three standardized experimental paradigms (**Fig. 1B-D**) in the macaques’ existing social settings to empirically assess group- and dyadic-level social relationships, tolerance (n = 105), and prosocial (n = 96) and cooperative tendencies (n = 102). Using these data, we evaluated within-group cooperative tendencies and investigated how variations in group-level outcomes might emerge from underlying patterns of dyadic cooperation.

**Fig. 1.**
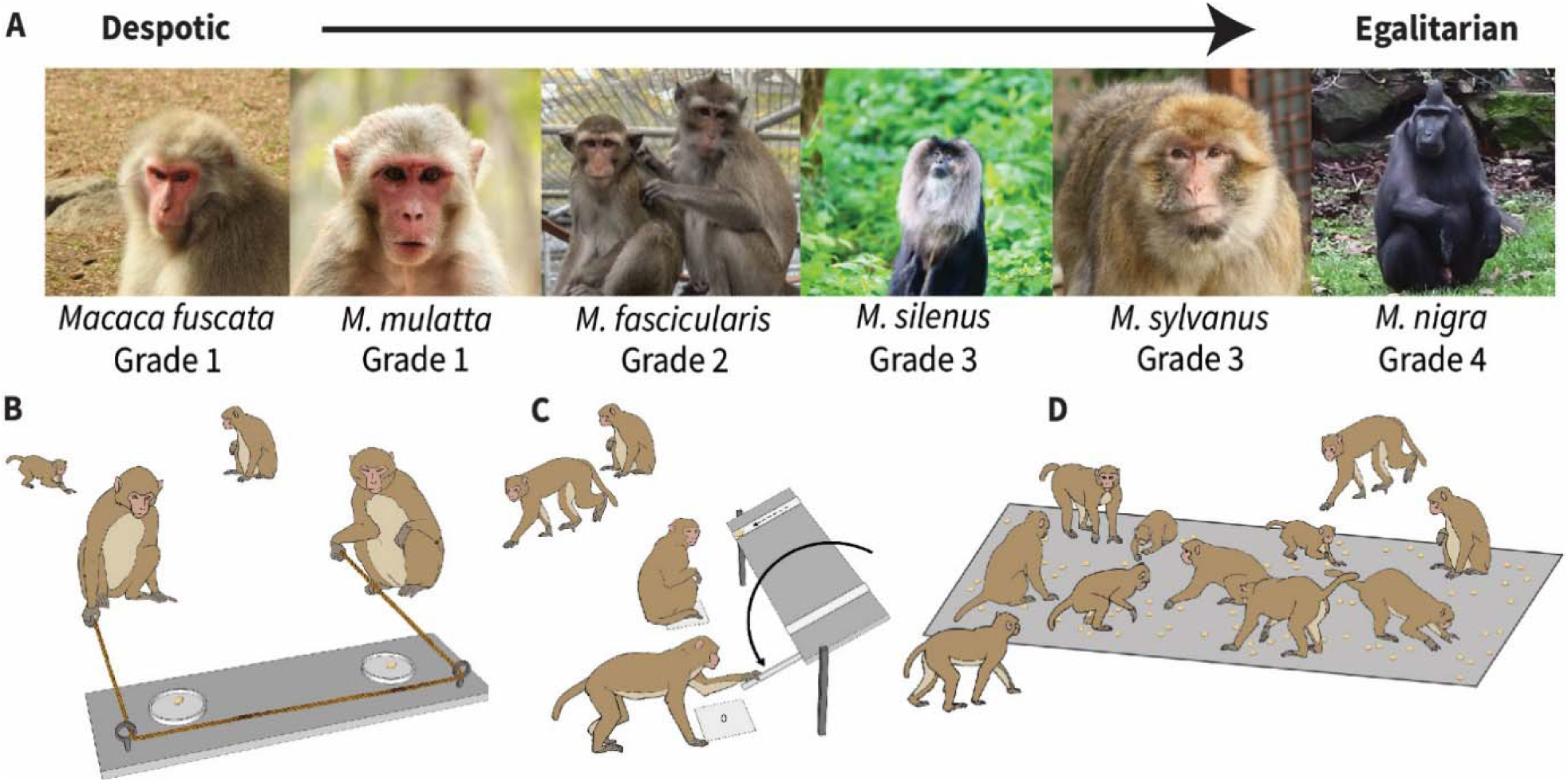
Study species of macaques and the different experimental paradigms. **(A)** Macaque societies are traditionally categorized into four grades based on their social styles. We studied six species representing all four grades – *Macaca fuscata* (Japanese macaque), *M. mulatta* (Rhesus macaque), *M. fascicularis* (Long-tailed macaque), *M. silenus* (Lion-tailed macaque), *M. sylvanus* (Barbary macaque), and *M. nigra* (Crested macaque). Due to macaques’ susceptibility to COVID-19, zoos and animal holding facilities implemented restrictive measures such as limited experimentation in close proximity. As a result, we could not collect data on all groups equally (**Table S1**), leading to different samples per experiment. We also used a small subset of our previously published data on cooperation (*12*) and prosociality (*13*, *27*) on single species for comparative analyses in the current study (cf. **Table S1**). **(B)** We tested macaques’ tendencies to cooperate using a loose-string paradigm (*10 groups and 102 individuals*, **Data S1**). Two self-trained individuals needed to pull two loose ends of a single string simultaneously to obtain food rewards (**Movie S1**). The string unthreaded when only one individual pulled, i.e., during uncoordinated pulling. We counted successful instances of cooperation without food monopolization out of 600 testing trials to calculate the group-level cooperation success rate. This experimental protocol also included a phase where dyadic social tolerance was investigated. **(C)** We used a group service paradigm to identify individuals with proactive prosocial preferences (*9 groups and 96 individuals*, **Data S2**). Individuals could proactively provide food to group members (**Movie S2**) by understanding the contingencies of two control (empty and blocked) conditions. We tested the tolerance of group members who participated in the group service experiment using a food distribution assessment phase (**Movie S3**). **(D)** We conducted a co-feeding tolerance test to assess the group-level tolerance of the macaque groups (*11 groups and 105 individuals*, **Data S3**). Contingent on group size, we created a ‘plot’ of a particular size where a predetermined number of peanuts were distributed (**Movie S4**). The cumulative proportion of a group in the plot at regular intervals or scans during eight experimental sessions provided information about their tolerance levels. Macaque photo credits (A): Jana Jäckels, Jorg Massen, Gaia Zoo, and Debottam Bhattacharjee. Illustration of experimental paradigms (B-D): Veera Schroderus.

## Results

### Macaque societies show within-group variation in cooperation and tolerance

Within-group cooperation success was assessed from the loose-string experiment (cf. **Fig. 1B, Data S4**), which ranged between 8.33% and 61.11% (mean ± standard deviation = 31.60% ± 18.75%). After controlling for group size effects, we found a strong negative correlation between cooperation success and macaque tolerance grades (Partial Bayesian correlation test (hereafter, correlation test): *r* = −0.85, *n* = 9, 89% credible interval or 89% *crl* = [−0.96, −0.54], **Fig. 2A**), suggesting that more despotic societies showed higher within-group cooperation success than more egalitarian societies. To understand the causes of this correlation, we first tested whether the macaque co-variation framework assumption regarding steepness of hierarchies (i.e., steeper hierarchies in more despotic societies) is met (cf. **Fig 1A**).

**Fig. 2.**
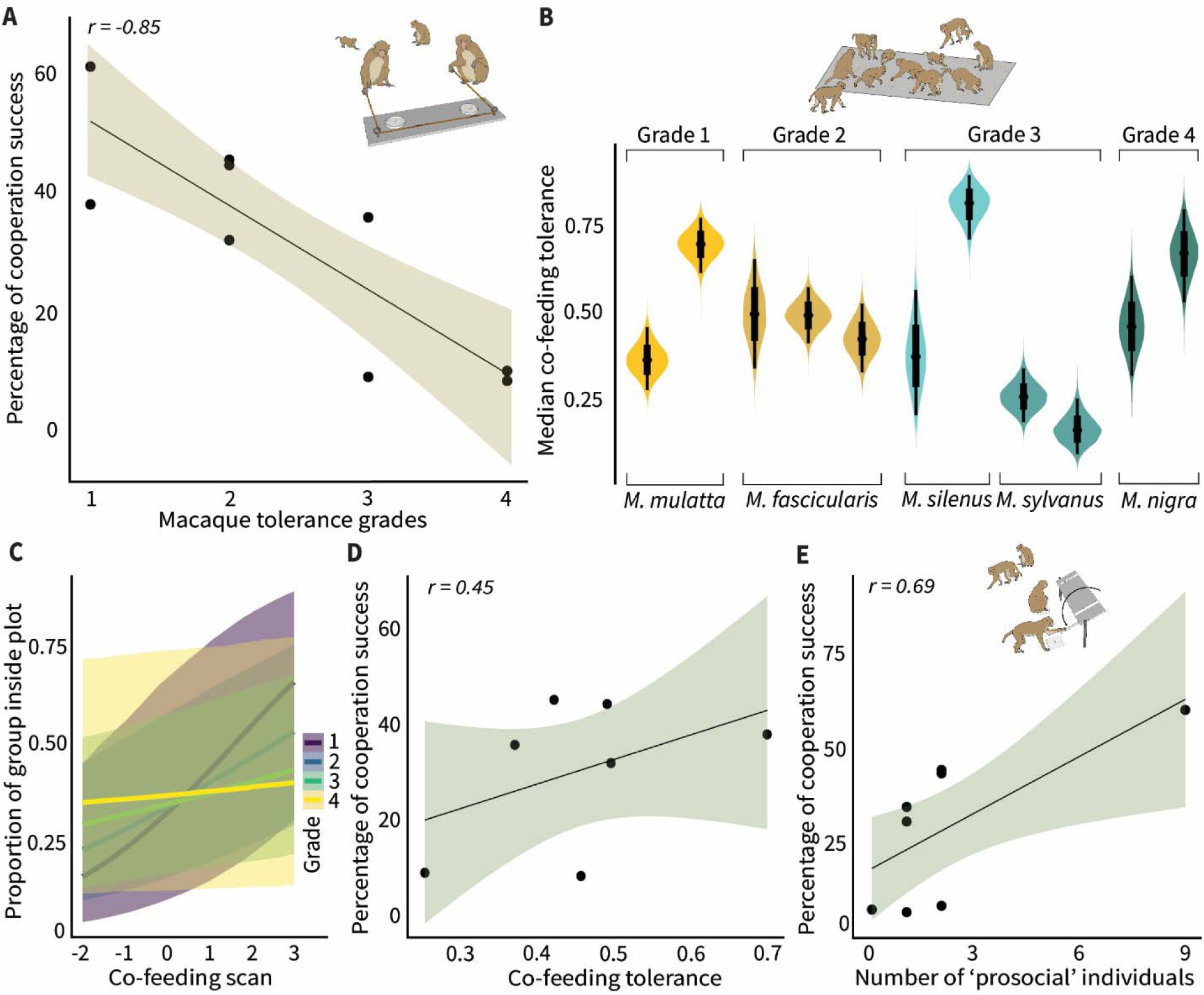
Within-group variation in cooperation and its associations with tolerance and prosociality. **(A)** Cooperation success in different species of macaques categorized under the four tolerance grades. A strong negative correlation indicates that more despotic macaque societies had higher cooperation success than egalitarian societies. **(B)** Intra- and inter-species variations in co-feeding tolerance levels. Co-feeding tolerance of the study groups belonging to five species and the four tolerance grades (**Data S3**). **(C)** The posterior predictive effects of an interaction between macaque tolerance grades and co-feeding session scans on the proportion of individuals of a group within the peanut plot, thus our proxy for co-feeding tolerance (**Data S3**). Colors represent the four tolerance grades. **(D)** Co-feeding tolerance and cooperation success had a weak positive correlation. **(E)** Higher cooperation success was associated with the presence of more individuals with prosocial motivations in groups. Solid dots indicate different macaque groups in all figures except B and C, and the shaded area shows an 89% credible interval. In B and C, the width of the violins represents data distribution (89% credible interval), and the solid black points indicate median values.

A negative but weak correlation was found between tolerance grades and hierarchy steepness (*r* = −0.33, *n* = 12, 89% *crl* = [−0.70, 0.19], **Data S4**), indicating a trend that despotic societies indeed have steeper hierarchies than egalitarian ones. However, the assumption of the co-variation framework is built upon within-group interactions among adult females (*18*, *19*, *21*). Accordingly, based on only adult females in the groups (applicable to 10 groups), we re-calculated steepness and re-investigated the relationship. In these data, a strong negative correlation was found (*r* = −0.61, *n* = 10, 89% *crl* = [−0.87, −0.11], **Data S4**), indicating that the assumption was clearly met. However, we concentrated on the full groups in the remainder of the analyses to optimize statistical power. This allowed all sex combinations to be included in investigations at the dyadic level, as between-sex bonds can also be important in primates (*28*, *29*). Next, we investigated within-group tolerance and prosociality and examined their associations with cooperation success.

Social tolerance is a central concept in primatology, where enhanced tolerance is positively linked to cooperation (*6*, *7*, *30*–*32*). Yet, varying definitions and methodologies are present concerning its ‘structural’ and ‘behavioral’ constructs (*33*). We assessed within-group social tolerance using a standardized co-feeding ‘peanut plot’ experiment (*34*–*36*) (cf. **Fig. 1D, Data S3**). We used ‘co-feeding tolerance’ to denote our experimentally obtained social tolerance. Substantial intra- and inter-species variation in co-feeding tolerance was found (**Fig. 2B**); however, more despotic societies showed a higher co-feeding tolerance tendency – reflected by the presence of a larger proportion of group within the plot – than egalitarian ones (**Fig. 2C**). We further conducted a correlation test between the co-feeding tolerance and Pielou’s evenness index (or Pielou’s *J*’ (*37*)), calculated from the food distribution assessment phase of the group service experiment (cf. **Fig. 1C**). A strong positive correlation was found between co-feeding tolerance and Pieulou’s *J*’ (*r* = 0.92, *n* = 6, 89% *crl* = [0.59, 0.99]), validating our co-feeding tolerance measure in capturing macaques’ within-group social tolerance. A low to moderately strong positive relationship was found between within-group co-feeding tolerance and cooperation success (r = 0.45, n = 7, 89% crl = [−0.30, 0.86], **Fig. 2D**). In addition, a strong positive correlation was found between within-group cooperation success and the number of individuals with prosocial motivations (or ‘prosocial’ individuals) those groups had (*r* = 0.69, *n* = 8, 89% *crl* = [0.14, 0.92], **Fig. 2E, Data S2**), corroborating the positive relationship between prosociality and cooperation (*7*, *31*).

### Prosociality, social tolerance, and kinship as predictors of dyadic cooperation

Of all potential dyads in Grade 1 to 4 societies, 31.9%, 36.9%, 46.1%, and 88.8% cooperated at least once, respectively (**Fig. 3A-D, Data S5**). To understand how the magnitude of cooperation (cooperation success ≥ 1) was distributed, i.e., whether fewer dyads cooperated more or all dyads cooperated uniformly, we performed normality tests (**Data S5**). More despotic societies (e.g., grades 1 and 2) showed positively skewed patterns, whereas normal patterns were observed in more egalitarian societies (e.g., grades 3 and 4) (**Fig. 3E-H**). The overall low percentages of cooperative dyads and the skewed distribution of magnitude suggest a higher dyad specificity, thus partner choice, in despotic than in egalitarian societies.

**Fig. 3.**
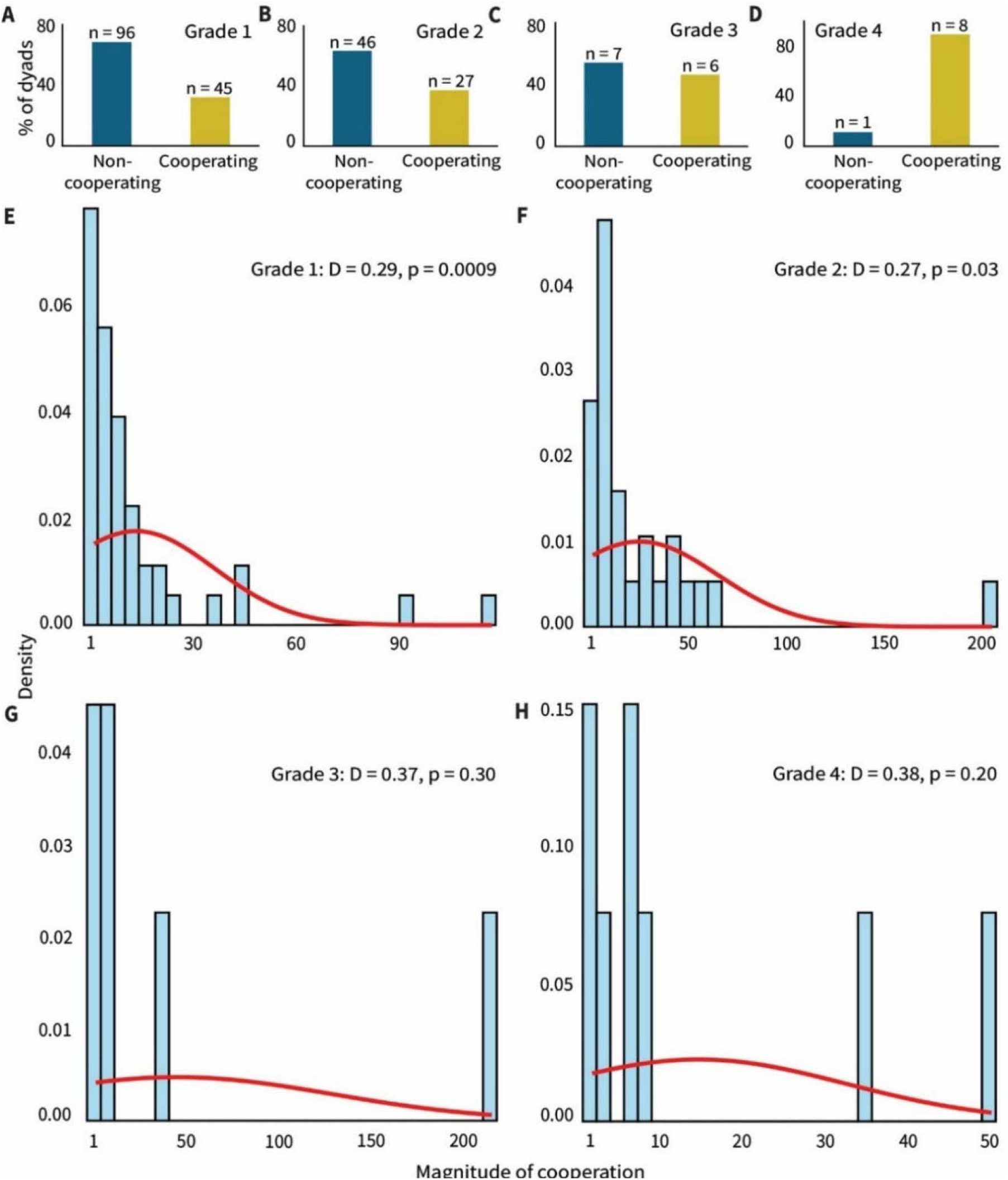
Dyad specificity in cooperation across macaque tolerance grades. **(A-D)** The percentages of cooperating and non-cooperating dyads in Grade 1, Grade 2, Grade 3, and Grade 4 societies. Note that the total number of potential dyads (i.e., both non-cooperating and cooperating) was calculated based on only self-trained individuals. **(E-H)** The distribution of magnitude values (cooperation success ≥ 1) in Grade 1, Grade 2, Grade 3, and Grade 4 societies with a histogram and an overlaid distribution curve (Kolmogorov-Smirnov test for normality with D statistic and p values). The histograms (in sky blue) represent the empirical distribution of the data, with the y-axes displaying the density of magnitude of cooperation. The red lines show the theoretical normal distribution based on the sample mean and standard deviation.

Due to the unavailability of uniform data across groups, two sets of models were built for analyzing the dyadic cooperation data (**Table S2-5**). In the first set of models on the dyadic cooperation data, we found evidence of prosociality, social tolerance, and kinship positively predicting the *likelihood* of cooperation (**Fig. 4A, Table S2**). Prosocial dyads were more likely to cooperate than non-prosocial dyads (*Est* = 1.41, 89% *crl* = [0.58, 2.27], *n* = 214, *pd* = 0.99). Dyads comprised of individuals with higher social tolerance were more likely to cooperate than dyads with lower tolerance (*Est* = 0.86, 89% *crl* = [0.40, 1.37], *n* = 214, *pd* = 0.99). Dyads with kin members were more likely to cooperate than dyads consisting of non-kin members (*Est* = 1.23, 89% *crl* = [0.29, 2.18], *n* = 214, *pd* = 0.98). Unlike in analyses on *likelihood*, only a strong positive effect of social tolerance was found in predicting the *magnitude* of cooperation (**Fig. 4B, Table S3**). Dyads comprised of individuals with higher social tolerance cooperated more than dyads with lower tolerance (*Est* = 0.23, 89% *crl* = [0.09, 0.40], *n* = 80, *pd* = 0.99). In the second set of models, except for a negative effect of group size on the *likelihood* (*Est* = −0.26, 89% *crl* = [−0.48, −0.07], *n* = 115, *pd* = 0.98), no effects of tolerance grades, grooming, and aggression indices were found on the *likelihood* and *magnitude* of cooperation (**Fig. 4C-D, Table S4-5**).

**Fig. 4.**
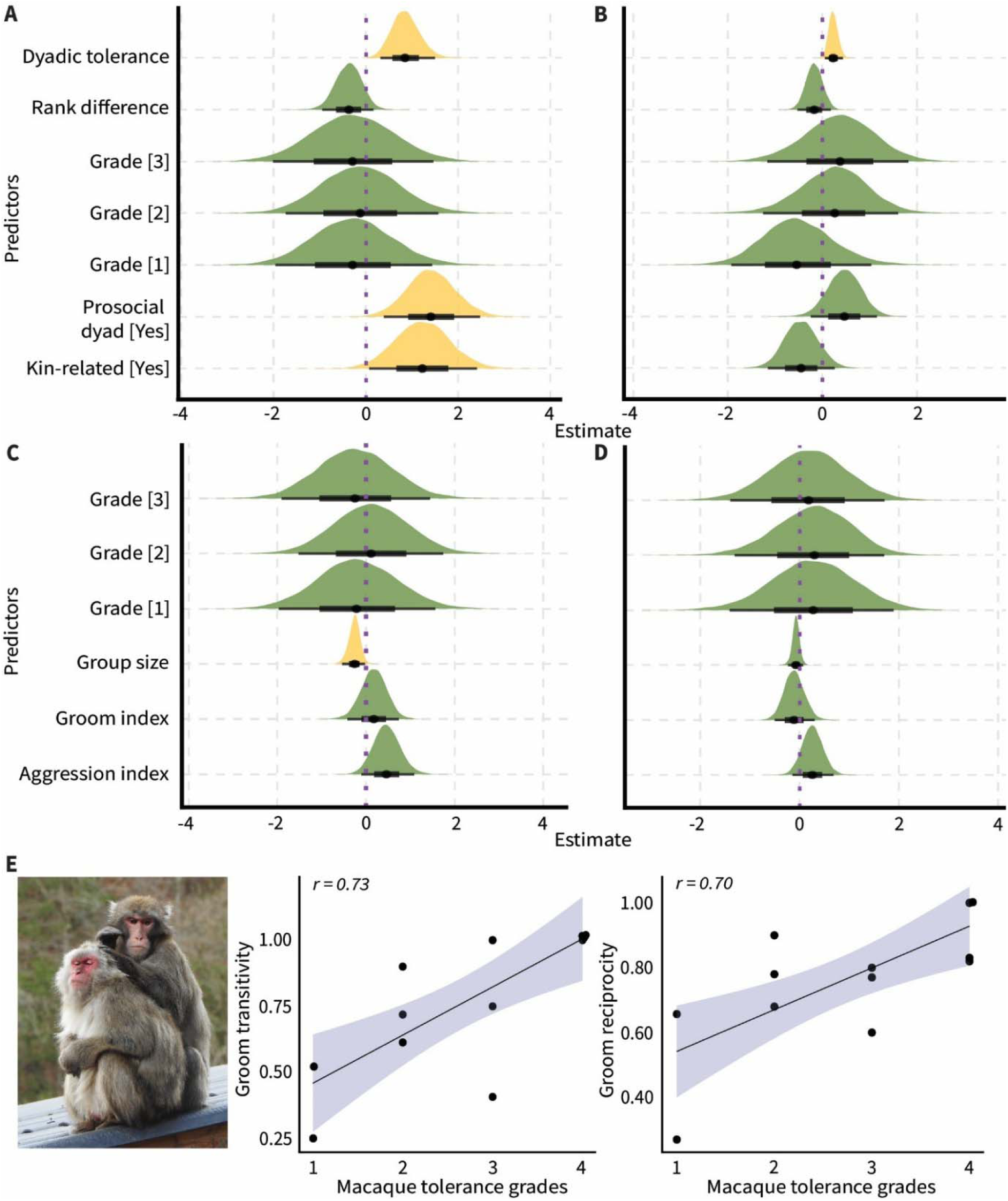
Dyadic cooperation and group-level grooming patterns. **(A)** The posterior predictive effects of dyadic social tolerance, rank difference, tolerance grades, prosociality, and kinship on the *likelihood* of cooperation. Yellow colors indicate robust effects of the corresponding predictors. **(B)** The effects of dyadic social tolerance, rank difference, tolerance grades, prosociality, and kinship on the *magnitude* of cooperation. Yellow color indicates a robust effect of the corresponding predictor. **(C)** The effects of tolerance grades, group size, grooming, and aggression indices on the likelihood of cooperation. **(D)** The effects of tolerance grades, group size, grooming, and aggression indices on the magnitude of cooperation. A and B represented results from a first set of models, whereas C and D represented results from a second set of models. **(E)** Group-level grooming transitivity and reciprocity showed strong positive correlations with macaque tolerance grades. Solid dots indicate the study groups, and shaded areas represent 89% credible intervals. In A-D, the width of the ‘half-eye’ represents data distribution (89% *crl*), and solid black points on the horizontal bars indicate median values. The vertical purple dashed lines indicate a parameter estimate of zero, i.e., the overlap of the credible interval with this line suggests no effect of the corresponding predictor. Macaque photo credit: Jana Jäckels.

In an additional set of Bayesian models, focusing on the results of the group-service prosociality experiment, we found that dyadic prosocial food provisioning was predicted by kinship and social tolerance (**fig. S2, Data S3, Table S6-7**). Kinship positively predicted both the likelihood (*Est* = 1.13, 89% *crl* = [0.33, 1.96], *n* = 302, *pd* = 0.99) and magnitude (*Est* = 0.98, 89% *crl* = [0.45, 1.51], *n* = 49, *pd* = 0.99) of prosocial food provisioning, whereas tolerance only positively predicted the likelihood (*Est* = 0.98, 89% *crl* = [0.57, 1.53], *n* = 302, *pd* = 1).

Results at the dyadic level corroborate findings that prosociality, social tolerance, and kinship are key proximate mechanisms of cooperation (*1*, *4*, *7*–*9*, *31*, *38*). Most importantly, they emphasize that cooperative dyads in more despotic societies are fewer and have higher selectivity in comparison to more egalitarian societies. We also expected this strong dyadic interdependency in despotic societies to be reflected in grooming interactions (cf. **Fig. 4C-D**). Therefore, we conducted additional analyses to understand within-group grooming patterns. Based on the frequency of grooming, we built social networks and calculated global transitivity, reciprocity, and modularity. Transitivity provides information on the level of clustering and is particularly useful for interpreting weighted and directed social networks (*22*, *39*). Reciprocity, by contrast, is a measure showing the difference between grooming efforts given and received. Modularity denotes how a network can be divided into communities where individuals groom specific partners more frequently than chance level (*23*). We found both transitivity and reciprocity to be strongly positively correlated with macaque tolerance grades (Transitivity: *r* = 0.73, *n* = 12, 89% *crl* = [0.38, 0.90]; Reciprocity: *r* = 0.70, *n* = 12, 89% *crl* = [0.33, 0.89], **Fig. 4E**), indicating the presence of more transitive and reciprocal grooming networks in more egalitarian societies.

Further, more egalitarian societies showed a lower tendency of grooming modularity than more despotic societies (*r* = −0.36, *n* = 12, 89% *crl* = [−0.72, 0.16]). Grooming patterns aligned with the assumptions of the co-variation framework (*18*, *19*, *21*). This opened up the idea that more despotic societies possibly restrict the number of strong (cooperative) bonds based on affiliative interactions (e.g., grooming) and that they may be used as a commodity traded in exchange for alternate services, like support during conflicts (*sensu* biological market theory) (*40*). To confirm, we constructed agent-based models simulating macaque social behavior along a despotic-egalitarian gradient.

### Emergence of strong bonds is restricted by dominance hierarchy in despotic societies

Strong social bonds can promote cooperation (*32*, *41*). To investigate how these bonds may emerge in despotic and egalitarian societies, we employed agent-based EMO-models (*42*–*44*). EMO-models simulate macaque social behavior, where ‘agents’ or individuals within a group are capable of emotional bookkeeping (*45*, *46*), integrating information of past affiliative interactions into a partner-specific LIKE attitude value. These LIKE relationships range between ‘0’ and ‘1’, with higher values indicating stronger bonds. To test the effect of despotism vs egalitarianism on LIKE relationships, we incorporated a *Hierarchy steepness* variable. Despotic, intermediate, and egalitarian societies were assigned steepness values of ‘1’, ‘0.6’, and ‘0.2’, respectively. The effect of steepness on LIKE relationships was investigated in diverse environmental conditions: three increase speeds (i.e., the speed with which LIKE attitude increases when grooming is received: fast, intermediate, and slow), three decrease speeds (i.e., the speed with which LIKE attitude decreases in the absence of being groomed; fast, intermediate, and slow), and two LIKE dynamics (‘easy-going dynamics’ and ‘picky dynamics’ (*47*)). In picky dynamics, it is more difficult to establish new relationships, but once a strong relationship is formed, the quality decreases more slowly over time compared to a relationship of the same quality in easy-going dynamics. We also assessed the stability of LIKE relationships over time (*47*). We investigated LIKE relationships at five different time points and calculated a correlation score between consecutive time points. The resulting four scores were averaged for each simulation to obtain a ‘stability’ score.

In line with our expectation, the number of potential interaction partners and the emergence of high LIKE relationships were restricted by the steep dominance hierarchy (values of ‘1’ and less so in ‘0.6’) in despotic societies (**Fig. 5A-C**). Strong bonds formed almost exclusively among individuals relatively close in their dominance ranks in more despotic societies. The pattern was consistent in situations of all decrease speeds. However, slower decrease speeds resulted in a higher absolute number of high LIKE relationships overall. In comparison to despotic and intermediate hierarchical societies, the dominance hierarchy had little influence on the emergence of high LIKE relationships in egalitarian societies (value of 0.2). When calculating the stability of the LIKE relationships, we found that they were more stable in despotic than egalitarian societies (**Fig. 5D**).

**Fig. 5.**
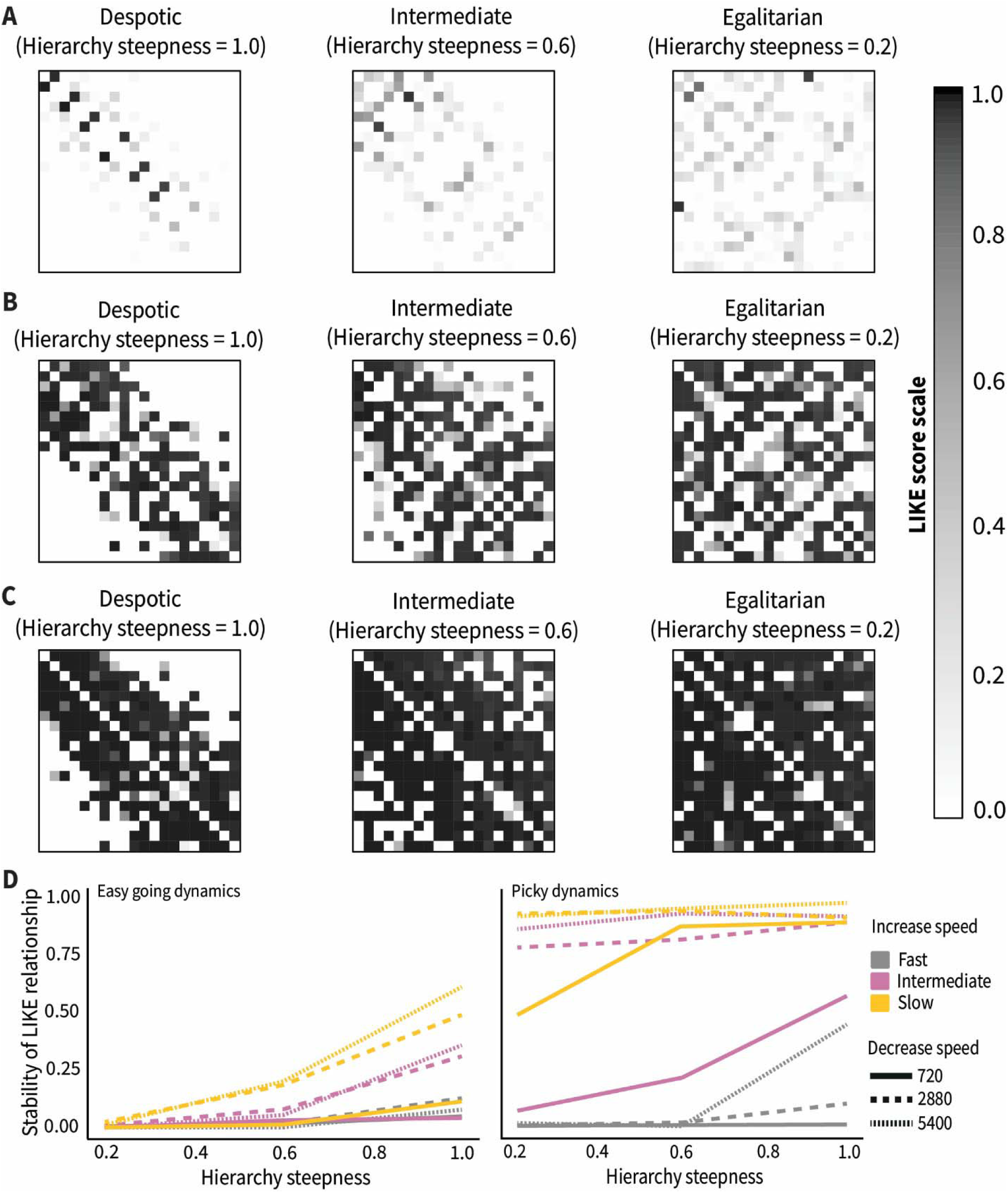
EMO-models show the emergence and stability of high LIKE relationships in societies along a despotic-egalitarian gradient. **(A)** LIKE relationships in societies with an intermediate increase speed, fast decrease speed, and picky dynamics. **(B)** LIKE relationships in societies with an intermediate increase speed, intermediate decrease speed, and picky dynamics. **(C)** LIKE relationships in societies with an intermediate increase speed, slow decrease speed, and picky dynamics. In A, B, and C, on the y-axes, individuals are ordered from low ranking (top row) to high ranking (bottom row), and on the x-axes, from low ranking (left) to high ranking (right). Each square represents a LIKE attitude from one individual to another, indicating their LIKE relationship. A LIKE relationship ranges from 0 (white) to 1 (black), with higher values indicating stronger bonds. **(D)** The stability of LIKE relationships in despotic (Hierarchy steepness = 1.0), intermediate (Hierarchy steepness = 0.6), and egalitarian (Hierarchy steepness = 0.2) societies using the easy-going and picky dynamics. Only one run is shown for all parameter settings. Repeats of runs using the same settings but a different random seed always resulted in very similar stability scores except for runs with picky dynamics, a slow increase speed, and a fast decrease speed. See **fig. S3-7** for EMO-model simulation runs with fast and slow increase speeds and easy-going dynamics. These simulations showed either similar results or very stable LIKE relationships across the whole population regardless of Hierarchy steepness (slow and intermediate increase and decrease speeds) or almost no LIKE relationships, regardless of the Hierarchy steepness (fast decrease and slow increase). As the latter two are not what we know from empirical data, these seem ecologically less relevant.

## DISCUSSION

Experimental evidence on the motivational mechanisms facilitating spontaneous cooperation was lacking, especially in less-tolerant societies (*48*). We show that more despotic societies exhibit higher within-group cooperation than egalitarian societies. Although evidence of a weak yet positive tolerance–corporation link was found, surprisingly, co-feeding tolerance did not align with the attributed social tolerance grades of macaque societies. This contrasting pattern could be due to including individuals of both sexes (and also juveniles) as opposed to social tolerance constructs that are built exclusively on interactions among adult females (*19*, *21*). Notably, species-level differences of co-varying traits in macaques along the despotic-egalitarian gradient are not always continuous (*23*, *49*). Social tolerance was also the best predictor of cooperation at the dyadic level, along with partial positive effects of prosociality and kinship. Our results, therefore, indicate selectively enhanced dyadic tolerance in more despotic societies, promoting cooperation.

In line with the prediction of the interdependency hypothesis, the percentages of cooperating dyads in despotic societies were lower, and the magnitude of success was even more restricted to specific dyads than in egalitarian societies. Agent-based models consistently showed that steeper hierarchies of more despotic societies allow for the emergence of fewer but, notably, more stable social bonds along the dominance hierarchy. Thus, steep hierarchies seem to force individuals into strong and interdependent bonds with those close in rank. Yet, based on our empirical data, the smaller rank differences only showed a tendency to promote cooperation positively (but see (*12*)). We suspect this overall effect might have been weakened by the data of more egalitarian societies that are not so restrictive regarding partner choice while cooperating. Nevertheless, this could also indicate that successful cooperative dyads in despotic societies do not exclusively rely on frequent reciprocal affiliative exchanges and instead may use ‘Machiavellian’ tactics (*50*, *51*), i.e., individuals may weigh costs and benefits for cooperative decision-making. Since hierarchies in macaque societies tend to be stable (*28*, *52*) (but see (*53*)), the relationships in despotic societies are too, and consequently more so than those within egalitarian societies. Therefore, selective strong bonds may enhance dyadic tolerance, ultimately facilitating (more) discriminate cooperation. These findings, together with the evidence of relatively higher asymmetric and lower transitive within-group grooming interactions, emphasize the role of strong interdependencies promoting cooperation in more despotic societies.

Interdependence – a key factor that shapes social tolerance – may vary across group- and dyadic levels. The emergence of dyadic cooperation was best predicted by social tolerance, even when controlled for the different species and groups. This aligns with the assumption of the interdependency hypothesis that strong dyadic interdependencies can result in higher tolerance, subsequently leading to higher cooperation (*31*, *32*, *54*–*56*). We also found the effects of prosocial motivations and kinship on the likelihood of cooperation, conforming with the findings of previous studies (*7*, *31*, *57*, *58*). Despite steeper hierarchies in more despotic societies, discriminate cooperation can be evolutionarily maintained through direct reciprocity and kinship (see discussion (*13*)). Moreover, individuals in more despotic societies can adopt ‘behavioral strategies,’ such as selective prosocial helping and cooperation (*12*, *13*), as rank-related fitness benefits are not always evident (*59*), and those strategies can potentially aid during active rank mobility (*53*).

In humans, hierarchies are known to influence cooperation, contingent on how they are operationalized. While a large dyadic rank difference can be detrimental to both human and non-human primate cooperation (*12*, *30*, *60*), ‘functionalist’ theories suggest that group or organizational level hierarchy promotes human cooperation through efficient decision-making, motivation, and coordination (see (*61*)). Although empirical evidence supporting the functionalist theories is mixed (cf. (*61*)), social relationships – through which dyadic interdependencies may form – are rarely investigated. Future comparative research incorporating social relationships and interdependencies may enable us to better understand the functional mechanisms of cooperation. In summary, our study challenges the conventional egalitarianism-cooperation view and provides rare and compelling evidence that cooperation can emerge and be sustained even in highly despotic societies through strong interdependencies.

## Materials and Methods

### 1. Study subjects, housing, and husbandry

We studied 13 captive groups of macaques belonging to six different species (*Macaca fuscata*, *M. mulatta*, *M. fascicularis*, *M. silenus*, *M. sylvanus*, and *M. nigra*) in their existing social settings. The *M. fuscata* group was housed at the Affenberg Landskron Park in Villach, Austria, in ca. 40,000 m^2^ enclosure with natural mixed forests typical to Southern Austria (*62*). The population size was ∼170. The two *M. mulatta* groups (R3G2 and R3G7) were housed at the Biomedical Primate Research Centre (BPRC) in Rijswijk, the Netherlands. The R3G2 and R3G7 groups comprised 30 and 25 individuals, respectively. Both groups had identically designed 74 m^2^ indoor and 250 m^2^ outdoor enclosures. The three *M. fascicularis* groups (J1G4, J1G7, and NWR) were also housed at the BPRC. The group sizes of J1G4, J1G7, and NWR were 15, 18, and 4, respectively. One individual was removed from J1G7 due to within-group compatibility issues before the completion of the studies and, hence, was not included in the analyses. J1G4 and J1G7 had identically sized 49 m^2^ indoor and 183 m^2^ outdoor enclosures. Due to a relatively small group size, the NWR group had 3.55 m^2^ and 3.88 m^2^ indoor and outdoor enclosures, respectively. We studied two *M. silenus* groups, one housed at Blijdorp Zoo and the other at the Apenheul Primate Park, both in the Netherlands. The Blijdorp and Apenheul groups consisted of 3 and 8 individuals, respectively. The Blijdorp *M. silenus* group had 100 m^2^ and 106 m^2^ indoor and outdoor enclosures, respectively, whereas the Apenheul *M. silenus* group had an 80 m^2^ indoor and a 768 m^2^ outdoor enclosure. Of the two *M. sylvanus* groups, one was housed at Gaia Zoo in the Netherlands and the other at the Apenheul Primate Park. The Gaia Zoo and Apenheul *M. sylvanus* groups comprised 14 and 13 individuals, respectively. The Gaia Zoo *M. sylvanus* group had a 108 m^2^ indoor (not for regular use of the animals) and a 3522 m^2^ outdoor enclosure. For the Apenheul *M. sylvanus* group, the only (outdoor) enclosure had an area of 3829 m^2^ with natural vegetation, rocks, and a narrow creek. Finally, the three *M. nigra* groups were housed at Bijdorp, Artis Zoo in the Netherlands, and Planckendael Zoo in Belgium, and the group sizes were 5, 4, and 6, respectively. Blijdorp *M. nigra* group had an indoor enclosure of 70 m^2^ and a 160 m^2^ outdoor enclosure. The Artis *M. nigra* group had 65 m^2^ and 761 m^2^ indoor and outdoor enclosures, respectively. The Planckendael *M. nigra* group had access to only an 82 m^2^ indoor enclosure. For groups with indoor and outdoor enclosures, connecting tunnels enabled macaques to move freely between them (except for the Gaia *M. sylvanus* group). The group sizes reported here included all individuals, including <1 year old. However, for our study, we only considered individuals >1 year at the time of testing (see **Table S1**).

All observational and experimental study procedures were approved by the Animal Experiments Committee and Animal Welfare Organisation of BPRC (Animal Welfare Organisation/IvD approval no.: 019A, 019C, 019D, and 019E). Furthermore, internal committees of all relevant zoos carefully monitored the study. All study components were non-invasive (European Directive 2010/63), and we strictly adhered to the ethical principles and guidelines of the American Society of Primatologists for the care and inclusion of animals. No animals were isolated from their existing social groups during our research.

The husbandry protocols differed slightly across study groups due to species-specific requirements and in-house management decisions, but these protocols adhered to the European Association of Zoos and Aquaria guidelines for accommodation and care for animals (*63*). Accordingly, all enclosures had multiple enrichment structures, like climbing platforms, hanging ropes, wooden structures, tree trunks, and slides. Except for the *M. fuscata* and Apenheul *M. sylvanus* groups, which live exclusively in outdoor enclosures, indoor enclosures of all groups were temperature-controlled and had concrete floors covered with sawdust bedding. Depending on the nutritional requirements, feeding routines also varied across groups. The diet primarily consisted of monkey pellets, fresh vegetables and fruits, and seed mix (e.g., sunflower and corn). All study groups had access to drinking water 24/7. No change in the regular feeding schedule was made for our study. The participation of individuals in all of our studies was completely voluntary. Since our study period overlapped with the COVID-19 pandemic years (overall study period: November 2021-February 2023), the zoos and animal holding facilities had strict preventive measures (e.g., restriction in behavioral experimentation in close proximity to animals) due to macaques’ susceptibility to COVID-19. As a result, we could not perform all the tests uniformly across our study groups. In addition, a small subset of data in this study has been used from published research on single species. See **Table S1** for details on the observations, different tests, the specific study groups on which we performed them, and the use of published datasets. Multiple experimenters were involved in the collection of data. However, the same experimenter(s) carried out all phases of a single test (cooperation, prosociality, and co-feeding tolerance) for a given study group to avoid potential experimenter bias. We used a randomized order in which behavioral observations, cooperation, and prosociality tests were conducted. The co-feeding tolerance test took place at the end for all study groups. Notably, the study groups were familiarized with the concerned experimenters and showed no distress during the investigation.

### 2. Cooperation test

We used a *loose-string paradigm* to assess the cooperative tendencies of macaques following a standardized protocol (*12*). In this paradigm, the experimental apparatus consisted of a movable platform (width = ca. 1.1 m) placed above a wooden base (width = ca. 1.3 m). Two plastic food trays were installed on either end along the side of the movable platform closest to the macaques. The other side of the platform had a handle attached, which an experimenter could use to move the platform. The food trays were far apart such that an individual could not simultaneously obtain food from both. Two small metal loops were anchored to the platform. Based on the experimental phase, strings were either attached or inserted through the loops, which allowed macaques to move the platform by pulling. As advised by in-house veterinarians, we used unshelled peanuts (halves) and mixed seeds (corn, sunflower, and pea) as food rewards for the cooperation experiment. We found that the participating individuals obtained all food rewards upon successful cooperation trials, suggesting their strong motivation for these food items. The cooperation test consisted of three phases: *habituation*, *training and social tolerance*, and *testing*, all of which were video recorded using Canon Legria HF G25 and Sony FDR AX100E cameras. During all phases, the identity of the individuals and how often they pulled the strings were recorded.

The habituation phase allowed macaques to familiarize themselves with the experimental apparatus. We placed the apparatus in front of the enclosure so the individuals could interact with it (through touching, licking, and sniffing). Multiple food rewards were placed on the food trays to get the attention of the individuals. The experimenter gently pushed the platform by using the handle for individuals to obtain food rewards. This step was repeated until 50% of the group members obtained at least three food items and became habituated. Habituating half of the *M. fuscata* group was practically challenging. Therefore, we adjusted the criterion for this group from 50% to >8% (i.e., at least more than 14 individuals) of the population. This percentage was chosen considering the sizes of other relatively large macaque groups in the study. The habituation phase was performed for a maximum of three hours on a single day, for 2-3 days, to fulfill the criterion and proceed to the next phase.

In the training and social tolerance phase, macaques voluntarily self-trained themselves with the pulling mechanism of the apparatus. Two small strings were tightly attached to the metal loops of the moving platform, such that pulling either string by an individual could bring the platform close to them. In addition to self-training, we used this phase to measure dyadic social tolerance among the participating group members. A trial began when the experimenter called “monkeys” and placed two similar food rewards on the two feeding trays. This procedure helped get the attention of the group members. After placing food rewards, the experimenter presented the strings to the macaques. We counted how often macaques obtained food rewards close to their (specific) group members at the other string. A trial concluded when both food rewards were obtained or after 2 minutes, whichever was earlier. If macaques did not pull the strings for three consecutive trials, the session was stopped and resumed the next day. A total of 18 sessions were conducted per group, each with 20 trials. Not more than four sessions were conducted on a single day. Approximately 20-second inter-trial and 5-minute inter-session intervals were used. Upon completion of sessions, we investigated whether 50% of each study group and >8% of the *M. fuscata* group obtained food rewards at least 10 times. All other groups except R3G7 (a rhesus group) fulfilled this criterion. We conducted five additional training sessions with R3G7, but the criterion was still not fulfilled. Nevertheless, we included this group in the next phase of testing.

The testing phase investigated macaques’ tendencies to cooperate with group members. A single and relatively long string was inserted through the metal loops and presented to the macaques. Thus, to move the platform and obtain food rewards, two individuals needed to pull the two loose ends of the string simultaneously. If only one individual pulled, the string unthreaded, and the platform did not move from its original position. A trial began when the experimenter placed food rewards on the trays, called “monkeys”, and presented the two loose ends of the string. A trial ended when two individuals simultaneously pulled the strings and successfully moved the platform or when the string unthreaded due to pulling by one individual. A trial lasted for a maximum duration of 2 minutes. We conducted a total of 30 sessions per group, with each session having 20 trials. The number of sessions conducted per day and inter-trial and inter-session intervals were identical to that of the training and social tolerance phase.

### 3. Prosociality test

We used a *group service paradigm* (*7*, *38*, *64*, *65*) to investigate the prosocial motivations of macaques following standardized protocols of seesaw (*13*) and swing set (*27*) mechanisms. While the seesaw mechanism was used predominantly in our study, COVID-19 restrictions forced us to use the swing set mechanism for the three *M. fascicularis* groups. This ensured that the macaques were tested from a relatively greater distance, unlike the seesaw mechanism. Notably, the two mechanisms were identical in their representations of the group service paradigm. We explained the seesaw mechanism below and highlighted its swing set counterparts.

The seesaw mechanism consisted of a wooden board (length = ca. 1.5 m) with two transparent pipes (diameter = ca. 3 inches) attached to two ends (Pos. 0 or provision end and Pos. 1 or receiving end). Food rewards (unshelled peanut, corn seed, sunflower seed, and pea seed) could freely move through the pipes. By default, the wooden board was tilted towards the experimenter’s end. A wooden or metal handle was connected to the board next to Pos. 0. The handle was projected towards the macaques’ end, which, upon pressing, could tilt the board towards them. When food rewards were present, pushing the handle resulted in items rolling down through the pipes and coming within reach of the macaques. However, an attempt to push or release the handle halfway resulted in the seesaw returning to its initial position, making food out of reach for the macaques. To obtain food rewards, it was thus essential to push the handle fully. Due to the distance between the food pipes, an individual pressing the handle (at Pos. 0) could not reach Pos. 1 to receive food rewards. Thus, macaques could only provision group members. The swing set mechanism used a 1.2 m long wooden beam hung from the ceiling by steel chains. Instead of food pipes, the apparatus included two small plastic buckets (at Pos. 0 and Pos. 1). Unlike the handle in the seesaw apparatus, a rope was attached next to Pos. 0, pulling which could bring the beam with food buckets close to the macaques, and again releasing it, e.g., to try and get access to the other bucket, resulted in the swing moving back to its original position in which the buckets were out of reach.

The group service paradigm consisted of the following phases: *habituation to apparatus*, *initial training and habituation to procedure*, *food distribution assessment*, *apparatus training*, *group service*, and *blocked control*. The first three phases included general habituation and self-training of the macaques. The prosociality test was video recorded using Canon Legria HF G25 and Sony FDR AX100E cameras. During all phases, the identity of the individuals and the number of times they obtained food rewards were recorded.

In the habituation phase, food items were spread over the wooden board (or placed inside the buckets in the swing set apparatus) at regular intervals for macaques to interact with the apparatus. This phase eliminated any potential effect of neophobia on the experimental setup. On consecutive days, we conducted a total of 2-3 sessions, with each session lasting 60-90 minutes. Individuals were considered habituated when they obtained at least 10 food rewards. Like the loose-string paradigm, we set a target of habituating at least 50% of group members of all study groups, except for the *M. fuscata* group, where this criterion was >8%. The criterion was fulfilled for all groups.

We decided the number of trials per session for the next phases by considering the sizes of the study groups. The number of trials was n*5 (n=group size with individuals >1 year) for all groups except *M. fuscata*. In the *M. fuscata* group, we found that 25 individuals were habituated; therefore, the number of trials per session was 25*5 = 125 (*13*).

The seesaw mechanism was fully locked during the initial training and procedure habituation phase, with the wooden platform tilted towards the macaques. Thus, macaques could not push the handle and operate the seesaw. However, upon placing in either position, macaques could retrieve food rewards that rolled down through food pipes. This initial training and habituation phase was not conducted for the swing set mechanism, as individuals were already habituated to the apparatus. Five sessions were conducted, and for each session, food rewards were placed in Pos. 0 and Pos. 1 alternately. A trial started when the experimenter called “monkeys” and placed a food reward in either Pos. 0 or Pos. 1. A trial concluded with an individual obtaining the reward or after a maximum of 2 minutes. Individuals were considered trained only if they obtained at least 10 food rewards. At least half of the habituated individuals needed to be trained to perform the next phase. This criterion was fulfilled for all groups.

The food distribution assessment investigated group-level social tolerance but only included individuals habituated to the apparatus. The procedure was identical to the previous phase, except the experimenter always placed food rewards in Pos. 1 in this phase. Two sessions were conducted to assess the average within-group food distribution or tolerance.

The seesaw (and swing set) apparatus was unlocked and operational in the apparatus training phase. Thus, macaques could push the handle (or pull the string to move the wooden platform/beam in the swing set). Unlike in the food assessment phase, the experimenter always placed food rewards in Pos. 0, such that the individuals operating the apparatus obtained rewards themselves. At least half of the habituated individuals needed to be trained by obtaining a minimum of 10 food rewards over five sessions. This criterion was fulfilled in all study groups except for two *M. fascicularis* groups (J1G4 and J1G7). Additional sessions (one for J1G4 and two for J1G7) were conducted (*27*), after which the criterion was fulfilled.

The group service phase consisted of test and empty control sessions. During test sessions, the experimenter placed food rewards in Pos. 1 of the apparatus. We recorded how often individuals pushed the handle (or pulled the string in the swing set) and provisioned food to group members. To control for the potential effect of stimulus enhancement (movement during food placement), we conducted empty control sessions, where the experimenter pretended to place food rewards in Pos. 1. A test trial started with the experimenter calling “monkeys” and placing a food reward in Pos. 1. The trial ended when an individual fully pushed the handle (or fully pulled the string in the swing set) or after 2 minutes. Note that pushing or pulling did not necessarily lead to food provisioning, as group members needed to be present at the receiving end (i.e., Pos. 1). Similar to the test sessions, an empty control trial began when the experimenter called “monkeys” and pretended to place a reward in Pos. 1, which ended when an individual pushed the handle or after 2 minutes. Five test and five empty control sessions were conducted alternatingly.

We blocked Pos. 1 using a plexiglass in the blocked control phase. This phase assessed whether pushing the handle (or pulling the string in the swing set) was due to self-motivation for food rewards. A trial began when the experimenter called “monkeys” and placed a food reward in Pos. 1. A trial concluded when an individual fully pushed the handle or after 2 minutes. Therefore, provisioning was impossible even if an individual was present at the receiving end (i.e., Pos. 1). Further, we conducted empty blocked control sessions where the experimenter pretended to place a reward in Pos. 1, similar to the group service phase. Alternatingly, five blocked and five empty blocked sessions were conducted. Note that we could not retrieve data from the first three blocked control sessions due to a technical malfunction with the video camera during the investigation of the R3G7 group (a rhesus group). However, to analyze whether individuals understood the paradigm and showed prosocial motivations, only the final two sessions (i.e., sessions 4 and 5) of each condition were considered (cf. (*7*, *13*, *38*, *64*, *65*)).

To keep the provisioning macaques motivated, all sessions of the group service and blocked control phases had motivational trials. During these trials, food rewards were placed in Pos. 0. Thus, provisioning individuals could obtain rewards by pushing the handle (or pulling the string in the swing set). Each session began with a motivational trial, repeated after every fifth regular trial. We used 10-second inter-trial and 5-minute inter-session intervals throughout the group service experiment. If macaques did not participate in three consecutive trials during a given session in a given phase (except empty and blocked control phases), the session was called off and resumed the next day. Besides, if ‘untrained’ individuals interacted with the apparatus during the group service or blocked control phase, we waited for them to move away, and repeated the affected trials. Notably, trials were conducted only when at least two trained individuals were present within a radius of 5 m of the experimental setup.

### 4. Co-feeding tolerance test

Based on the group sizes (considering individuals >1 year), we created testing zones or plots in macaques’ enclosures. The width of the plot was always 75 cm, and the length depended on the group size ((group size – 1) x 12.5 cm). For example, in a group of 5 individuals over one year, the plot length was 12.5 x 4 = 50 cm. We uniformly distributed a predetermined number of unshelled and unsalted peanuts (group size x 12 pieces) within the plot. The fundamental assumption was that, in a ‘tolerant’ group, all individuals would remain in close proximity to each other in a zone full of valuable resources. The proportion of a group present in the plot was used as a proxy for co-feeding tolerance, with a higher proportion suggesting greater tolerance.

The co-feeding tolerance test was conducted primarily in the indoor enclosures of the study groups. However, the test was performed in outdoor enclosures for the two *M. sylvanus* groups and one *M. silenus* group (in Apenheul) due to logistic advantages. Note that the *M. sylvanus* group in Apenheul had only a usable outdoor enclosure. We chose strategic areas in the enclosures that are typically used for feeding. These areas were chosen to make the plots, eliminating the bias of ‘not finding’ them by macaques. We made visible boundaries in the outdoor enclosures, and the designated areas were cleared of sawdust bedding for indoor enclosures, making them clearly visible. Groups were temporarily locked in their outdoor enclosures (indoor for Gaia *M. sylvanus* group and Apenheul *M. silenus* group). The experimenter prepared the plot in the designated areas using measuring tapes and uniformly distributed the predetermined number of peanuts. Our study groups receive enrichment food after general lock-unlock procedures (e.g., during enclosure cleaning) as part of standard husbandry protocols. Therefore, we expected the macaques to be motivated to visit the indoor enclosures (and not remain in outdoor enclosures). The experimental design differed slightly for the Apenheul *M. sylvanus* group. As per veterinary advice, we used hazelnuts instead of peanuts throughout all the sessions. Besides, since this group had only an outdoor enclosure, we did not have the opportunity to lock them elsewhere temporarily. Instead, after preparing the plot, hazelnuts were quickly distributed out of sight of the macaques, followed by catching their attention.

A total of eight sessions were conducted in each study group, except for the *M. nigra* group housed at Planckendael, where we could conduct seven sessions due to veterinary regulations. We controlled for this variation in our statistical analysis. Not more than one session was carried out on a given day. To avoid any potential bias of hunger, co-feeding test sessions took place at least one hour after the regular feeding schedule. A session started when an individual (or individuals) entered the plot and concluded when all the peanuts were eaten. We recorded the activities of the macaques for an additional 2 minutes after the plot became empty. The co-feeding tolerance test was video recorded using Canon Legria HF G25 and Sony FDR AX100E cameras mounted on tripods.

### 5. Behavioral data collection using continuous focal sampling

Behavioral observations were conducted using continuous focal sampling methods. We conducted 20-minute long focal sessions, and after correcting for the time out of sight, we obtained >29,800 minutes of behavioral data (**Table S1**). The observation minutes per group varied (cf. **Table S1**). The focal observational data were used to investigate the dominance-rank relationships, as well as direct affiliative and aggressive interactions. These measures were calculated at the group as well as dyadic levels. Focal data on the *M. fuscata* group could not be collected (cf. **Table S1**), but other ongoing and published research on the *M. fuscata* group provided us with information on dominance-rank relationships (*13*, *62*). Complete data on individual kin relationships were available for all study groups.

We utilized five behavioral variables representing unprovoked submissive behaviors (*avoid*, *be displaced*, *silent-bared teeth*, *flee*, and *social presence*) to construct within-group dominance hierarchies and to assess dyadic dominance-rank relationships (*66*). We recorded the frequency of grooming given and received to assess direct affiliative interactions. In addition, we collected data on proximity (within one body length) and affiliative body contact, which were used to validate the grooming data. Finally, we used seven behavioral variables representing contact- and non-contact aggression (*hit*, *chase*, *lunge*, *bite*, *grab*, *open-mouth threat*, and *stare*) to determine the aggressive interactions (*67*).

### 6. Data coding and statistical analyses

All data were coded from the video recordings using BORIS (version 7.13.6) (*68*) or in a frame-by-frame manner using Pot player (version 240618). Due to data coding by multiple coders, we comprehensively checked for inter-rater reliability using intra-class correlation (ICC) tests among the trained coders. Overall agreement was high, with the ICC (3,k) ranging between 0.88 and 0.97. Statistical analyses were performed using R (version 4.4.1) (*69*). The agent-based EMO-model was constructed using NetLogo (version 5.3.1).

We applied a Bayesian mixed modeling approach to analyze our empirical data using the *brms* package of R (*70*). We used weakly informative priors to guide the estimation process while allowing the data to dominate the posterior distributions (*71*, *72*). A normal prior with a mean (m) of 0 and a standard deviation (SD) of 5 was used for the intercept term. We applied normal priors for all other fixed-effect coefficients with m = 0 and SD = 1. An exponential prior with a rate parameter of 1 was assigned to the standard deviation of the random effects. Unlike *p*-values in the frequentist approach, we reported median estimate coefficients (*Est*), 89% credible interval (*crl*) that contains 89% of the posterior probability density function, and probability of direction (*pd*), indicating the direction and certainty of an effect. Model convergence was assessed following Bayesian statistics guidelines (*73*). We investigated trace and autocorrelation plots, Gelman-Rubin convergence estimations, and density histograms of posterior distributions of all the models. We sampled posterior distributions using 10,000 iterations and 2000 warmups.

#### Cooperation

We defined cooperation (or cooperation success) as two individuals in the testing phase simultaneously pulling the strings and obtaining rewards without monopolization. Our analyses focused only on the participating individuals (i.e., at least one pull) who were fully habituated and trained following the different criteria. The percentage of participating individuals varied across groups. We found extremely low participation in R3G7 (one of the *M.mulatta* groups). Notably, this is the same group in which we conducted additional training sessions. We decided to discard the data on this group from cooperation analyses. The range of group-level participation rate was 35.71% to 100% (mean = 66.94%, standard deviation = 20.05%), excluding the semi free-ranging *M. fuscata* group, where the participation rate was 9.41%.

Cooperation success was coded as a binary variable at the trial level. We counted how often (i.e., in how many trials) two individuals successfully cooperated in a group (see **Data S1** for the list of trained individuals). The total number of trials in the testing phase for each group was 600. However, the removal of an individual from J1G7 (one of the *M. fascicularis* groups) led us to adjust the number of trials in which the individual participated (cf. (*12*)). This resulted in an adjusted 459 testing trials in J1G7. We incorporated an ‘offset’ term in our statistical models to control the varying numbers of testing trials. To assess whether macaques generally understood the need for partners to obtain food rewards, we compared the instances when they pulled strings alone vs. in the presence of partners. We found that macaques pulled the strings more often with a partner than when present alone (Wilcoxon signed rank test: *Z* = −2.18, Cohen’s d = 0.59, 95% CI [0.06, 1.08], *P* = 0.028). Although delayed control trials may provide a better understanding of the cognitive processes underlying the need for cooperation partners (*74*), our measure in a social experimental setting potentially indicated the same (also see (*12*)). Additionally, we were more interested in investigating the spontaneous collaborative efforts of two individuals rather than the cognitive mechanisms facilitating active partner recruitment.

Data from the social tolerance phase of cooperation test was used to measure dyadic tolerance levels. The social tolerance phase allowed individuals to co-feed or monopolize the food rewards. We followed a standardized method, where all co-feeding and monopolization opportunities were considered (*12*). The number of times two specific individuals obtained food by sitting in proximity (i.e., two attached strings at the apparatus) was divided by the sum of these individuals feeding alone and monopolizing. The total number of trials in the social tolerance phase was 360 for each group, except for J1G7. Due to the removal of an individual in J1G7, similar to the testing trials, the number of social tolerance trials was adjusted. Instead of 360, it was 192 for J1G7. For each study group, dyadic social tolerance values were standardized, with higher values indicating higher dyadic social tolerance.

We investigated the effects of eight social and demographic dyadic attributes or predictors – prosocial motivations, dyadic social tolerance, kinship, rank- and age differences, sex compositions, and dyadic affiliative and aggressive relationships on cooperation. Empirical evidence suggests that dyads composed of at least one individual with prosocial motivation can lead to higher cooperation success than dyads consisting of individuals without any prosocial motivations (*31*). Accordingly, we called a dyad ‘prosocial’ when at least one of the members was identified to have some prosocial motivation (**Data S2**). For the dyadic social tolerance measure, we analyzed data obtained from the dedicated social tolerance phase of the loose-string paradigm (cf. **Fig. 1B**) (*12*). Dyadic social tolerance was measured by the tendency of individuals to obtain food rewards – with and without monopolization – in the presence of group members. We used a genetic relatedness cutoff of 0.25 to determine whether a dyad was kin-related. When calculating the within-group steepness of hierarchies (*75*), we determined the dominance-rank relationships, particularly the ordinal rank differences, among individuals. Focal data were used to create dyadic grooming and aggressive indices (*76*).

Theoretically, each self-trained participating individual within a group (cf. **Data S1**) had the opportunity to cooperate with other participating members. Subsequently, we used a sequential decision-making hurdle approach to analyze our zero-inflated cooperation data. In this approach, we first investigated the probability or likelihood of cooperation (first hurdle: yes or no, i.e., whether a dyad successfully cooperated at all), followed by the intensity or magnitude of cooperation (second hurdle: dyadic cooperation success ≥ 1). We used two sets (each encompassing eight groups) of Bayesian mixed models (i.e., hurdle models) for the analyses as data on socio-demographic factors based on behavioral observations were not uniformly available for the groups (cf. **Table S1**). The first set of models investigated the likelihood and magnitude of cooperation with all dyadic social and demographic attributes, except for grooming and aggression indices. These two indices were used as predictors in a second set of models. The relevant models included rank difference and grooming index as interaction terms with the tolerance grades. Interaction terms were dropped in case no robust effect was found in subsequent simpler models. Individuals nested within groups and groups nested within species were included as random effects in the models.

#### Prosociality

Participation (i.e., with complete habituation and training) in the group service paradigm was slightly higher than in the cooperative loose-string paradigm. It ranged between 61.50% and 100% (mean = 81.48%, standard deviation = 12.98%), excluding the *M. fuscata* group, where the rate was 14.70% (cf. (*13*)). Subsequently, instead of using the percentage of food provisioning (cf. (*7*, *64*)), we reported the number of individuals with prosocial motivations in a group as a measure of group-level prosociality.

We used three criteria to determine whether an individual had prosocial motivation. First, an individual needed to push the handle (or pull the rope in swing set apparatus) more often in group service test than at least one of the control conditions (i.e., empty control and blocked control). This criterion was set because the number of test trials in which individuals could push the handle was restricted by the number of participants and the number of pushes by other individuals. Therefore, pushing significantly more in the test than in both empty and blocked control conditions was a strict and conservative cut-off. Second, although pushing the handle and subsequently provisioning food may require spatiotemporal coordination among group members, i.e., the presence of a receiver at Pos. 1, an individual was only considered to have prosocial motivations when food was provisioned to a group member. Third, an individual proactively provisioned food to group members, i.e., without relying on solicitation from the receiver or showing aggression towards the receiver (*13*, *27*, *64*). We used Fisher’s exact tests to investigate whether an individual pushed the handle more in the test than at least one of the control conditions. Notably, we only used data from sessions 4 and 5 (of group service test, empty control, and blocked control) for analyses as macaques needed to learn the contingencies of the control conditions (*7*, *13*, *27*, *38*, *65*). The number of individuals with prosocial motivations ranged between 0 and 9 across groups (mean = 2.33, standard deviation = 2.59, **Data S2**). All individuals, other than the ones with prosocial motivations (i.e., with experimental evidence), were considered ‘non-prosocial’. This was decided as all group members had the opportunity to participate and provide food (if prosocial) to each other in the group service paradigm.

The food distribution assessment phase provided information on group-level tolerance but was restricted to only individuals who interacted with the apparatus. We calculated Pielou’s evenness index (or Pielou’s *J*’) (*37*). The index can range between 0 and 1, with higher values (i.e., closer to 1) indicating more uniform food distribution among group members. In our study groups, Pielou’s *J*’ ranged between 0.13 to 0.65 (mean = 0.41, standard deviation = 0.19, **Data S4**).

#### Co-feeding tolerance

For estimating group-level co-feeding tolerance, we calculated cumulative presence, defined by the proportion of individuals in the group present both inside the plot and within one arm’s length of reach of the plot at regular time intervals. To eliminate any potential effects of varying enclosure sizes and distances between unlocking zones (if applicable) and the plot, counting started (i.e., first scan) when an individual first entered the plot. Scans were conducted at each 10-second interval until the plot became empty. The number of scans per session per group varied greatly (range = 2 – 27, mean = 8.15, standard deviation = 4.91), primarily depending on the level of participation and monopolization rate. That is, more monopolization led to longer feeding duration, resulting in a higher number of scans. By investigating the obtained mean value, we decided on a cut-off of eight scans consistently across all sessions for all groups. Thus, data collected up to eight scans were utilized for the co-feeding tolerance analyses. During statistical analysis for the co-feeding tolerance, we also checked the results with an even smaller cut-off of six scans for validation. Macaque tolerance grades (i.e., grades 1 to 4) were included as an interaction term with scans. We constructed an intercept-only Bayesian mixed model to obtain the median co-feeding tolerance values of the groups. The median co-feeding tolerance ranged between 0.15 and 0.81, with higher values suggesting a higher co-feeding tolerance. Additionally, statistical models were built to compare macaque societies along the despotic-egalitarian gradient, where the cumulative presence variable was used as a function of the categorical tolerance grades and scans.

#### Dominance-rank relationships

We calculated group-level hierarchy steepness and determined the ordinal ranks of individuals along the corresponding hierarchies. A Bayesian Elo-rating method was used for these calculations (*75*). Based on unprovoked submissive behaviors, we prepared directional matrices. The Bayesian Elo-rating method calculates winning probabilities from these matrices to assess the steepness of the hierarchy. A steepness range of 0.23 to 0.91 was found (mean = 0.65, standard deviation = 0.22, **Data S4**), with higher values indicating more steeper hierarchies. For each group, dyadic rank differences were calculated and standardized for statistical analysis. As the macaque co-variation framework is built upon interactions among adult females, we also calculated within-group hierarchy steepness based on only adult females. Using these data, we found a steepness range of 0.24 to 0.93 (mean = 0.61, standard deviation = 0.24, **Data S4**).

#### Affiliative and aggressive interactions

We carried out social network analyses to measure within-group affiliative interactions. Specifically, we calculated global transitivity, reciprocity, and modularity values from grooming matrices (**Data S4**). In more egalitarian macaque societies, grooming distribution is less likely to be asymmetric, as less steep hierarchies in egalitarian societies do not hinder grooming from being symmetrically present (*19*, *21*, *22*). In contrast, in more despotic societies, most grooming interactions are concentrated among a few group members (typically including the dominant individuals or kin relatives), making the grooming networks modular and asymmetric. Within-group grooming transitivity values ranged between 0.24 and 1 (mean = 0.76, standard deviation = 0.26), with more transitive networks indicating a symmetrical distribution of grooming among group members. Reciprocity ranged between 0.27 to 1 (mean = 0.75, standard deviation = 0.19), with higher values suggesting more symmetrical or reciprocal grooming patterns. Finally, modularity had a range of 0 to 0.33 (mean = 0.76, standard deviation = 0.26), with higher values indicating the presence of more distinct modular grooming networks.

We calculated the group-level frequencies of aggressive interactions. All instances of aggressive behaviors were combined and corrected for the group’s overall observation minutes to obtain within-group aggression values. Notably, tolerance grades did not correlate with the frequency of aggression per minute (*r* = 0.14, *n* = 12, 89% *crl* = [−0.37, 0.59], **Fig. 2B**), suggesting that the frequency of aggression was comparable across the despotic-egalitarian gradient.

In addition to group-level measures, we calculated dyadic affiliative and aggressive indices using standardized methods (*12*, *76*). The observed frequencies of grooming, proximity, body contact, and aggression were extracted separately for all combinations of dyads. These values were then divided by the combined observation minutes of the individuals making the dyad. The resulting rates were further divided by a group average of the corresponding behavior (i.e., grooming, proximity, and body contact). Finally, we z-transformed the values to control for group-level variance. We specifically used grooming data for our analyses. To see whether grooming effectively captured affiliative relationships, we carried out Bayesian correlation tests between dyadic grooming and dyadic proximity (*r* = 0.45, *n* = 203, 89% *crl* = [0.34, 0.55], *pd* = 1), and dyadic grooming and affiliative body contact (*r* = 0.10, *n* = 203, 89% *crl* = [−0.04, 0.22], *pd* = 0.92). These results indicate that dyadic affiliative interactions, to a certain extent, were positively reflected by grooming.

#### Agent-based EMO-model

The agent-based EMO-model simulates macaque social behavior, where all ‘agents’ or individuals share the same behavioral rules but have variable internal states (*42*–*44*, *47*). These rules and internal states determine how actors interact, further influencing their internal states. Besides moving around in space, agents can engage in affiliative (grooming and affiliative signaling) and agonistic (attacking, aggressive, or submissive signaling) behavior. The EMO-model has been validated by comparing its results to empirical data on various species of free-living macaques (*44*).

The agents in the EMO-model are capable of emotional bookkeeping, integrating information of past affiliative interactions into a partner-specific LIKE attitude, and establishing a LIKE relationship. More frequent affiliative interactions (i.e., receiving grooming from a group member) increase the LIKE attitude associated with the group member, positively affecting the likelihood of affiliative social interactions with this group member (*44*). Reciprocal grooming relationships emerge in the model when grooming and LIKE within a dyad are mutually reinforcing. However, this only occurs in a limited part of the parameter space, provided that actors are very selective when choosing a partner (*42*) and that LIKE attitudes decrease over time with intermediate speed (*43*). However, the decreased speed that leads to the most naturalistic scenario, in turn, depends on the speed with which LIKE increases during grooming and the dynamics with which it increases and decreases (*47*). In contrast to a LIKE attitude, every agent also assigned a FEAR attitude to every other agent. The FEAR attitude does not change over time or due to social interactions; thus, it models a static dominance hierarchy. For instance, the most dominant individual (individual A) has a FEAR value of 1/20, while the least dominant individual has a FEAR value 20/20. FEAR-attitudes (from individual A towards individual B) are calculated by subtracting individual B’s FEAR value from individual A’s FEAR value (in this example: 0.05-1 = −0.95). FEAR attitudes, therefore, range from −0.95 (A is dominant over B) to 0.95 (B is dominant over A).

In order to manipulate the steepness of the dominance hierarchy, we added a variable to the EMO-model called *Hierarchy steepness*. The Hierarchy steepness variable ranged from 0.2 to 1, with higher values indicating more despotic societies (‘1’ = despotic, ‘0.6’ = intermediate, and ‘0.2’ = egalitarian). This variable scaled the range of FEAR attitudes in a group. For example, in a simulation run with a Hierarchy steepness setting of 0.1, FEAR attitudes would only range from −0.095 to 0.095, making the hierarchical (or rank) difference between the most dominant and least dominant individual smaller than 0.1. Since FEAR attitudes only impact the likelihood of aggressive and submissive behavior, the manipulation of hierarchy steepness only directly impacts the likelihood of aggression and submission. In contrast, the likelihood of affiliative behavior was only affected indirectly. To study the impact of Hierarchy steepness on the distribution of social relationships (as measured by LIKE values), we performed every simulation run three times, with each repeat having a different setting of Hierarchy steepness. The effect of Hierarchy steepness was investigated for two different dynamics settings (the original or easy-going, following (*42*–*44*) as well as the alternative or picky, following (*47*)), three different increase speeds (fast, intermediate, and slow; cf. (*47*)) and three different decrease speeds (720, 2880 and 5400, cf. (*43*, *47*)). The easy-going LIKE dynamics settings assume that increasing LIKE follows a linear curve while decreasing LIKE follows an exponential curve. In contrast, LIKE increases and decreases following a logistic curve while using the picky dynamics settings. As a result, when using the picky dynamics, it is more difficult to form new strong relationships, but once a strong relationship is established, its quality decreases more slowly over time, compared to a relationship of the same quality in a simulation run using the easy-going dynamics setting. All simulation runs were performed with very high partner selectivity (0.99; cf. (*42*)).

We investigated the impact of Hierarchy steepness on the distributions of LIKE values (averaged over the ‘second year’ of the recording period). Level plots were made for each simulation run, showing the LIKE attitude of agents toward each other (resulting in 20 x 20 level plots). Level plots were visually inspected. To quantify the stability of LIKE over time, LIKE distributions were measured at five different points in time, separated by 0.25 years (start of the second year of the recording period, after 1.25, 1.5, 1.75 years, and at the end of the recording period). For stability assessment, an R2 was calculated for each transition (i.e., between LIKE distributions of consecutive time points). The resulting four R2 values were averaged, resulting in an average ‘stability’ score for each simulation run (see (*42*–*44*, *47*) for details). Notably, all group-level properties in the EMO-model were emergent properties determined by the interactions among the agents.

Every unique run (i.e., every set of parameter settings tested) was repeated three times. These runs with the same parameter settings, but using a different random seed, always resulted in very similar stability scores. This was true for the entire parameter space except for runs with picky dynamics, a slow increase speed, and a fast decrease speed. For these runs, a fourth repeat was performed, to get a better understanding of what the more likely outcome would be. Although it still generally holds true for these settings that a steeper hierarchy will restrict the LIKE distribution, resulting in a more stable LIKE distribution, this is not true for every repeat. Because these runs have a slow increase speed and a fast decrease speed, combined with picky dynamics, it is very difficult to form a high-LIKE relationship, especially with a very steep hierarchy. This results in a scenario with only 1 or 2 high-LIKE dyads that remain stable over time, resulting in a high stability score. However, in some instances, there are no dyads that managed to form a high-LIKE relationship, resulting in a scenario without any stable high-LIKE relationships and a correspondingly low stability score (of 0). When hierarchy strength is low enough (Hierarchy strength = 0.2), it seems slightly easier to form somewhat stable relationships. While these runs can also be very stable, similar to the more despotic runs, no stability scores under 0.3 were observed.

---

**fig. S2.**
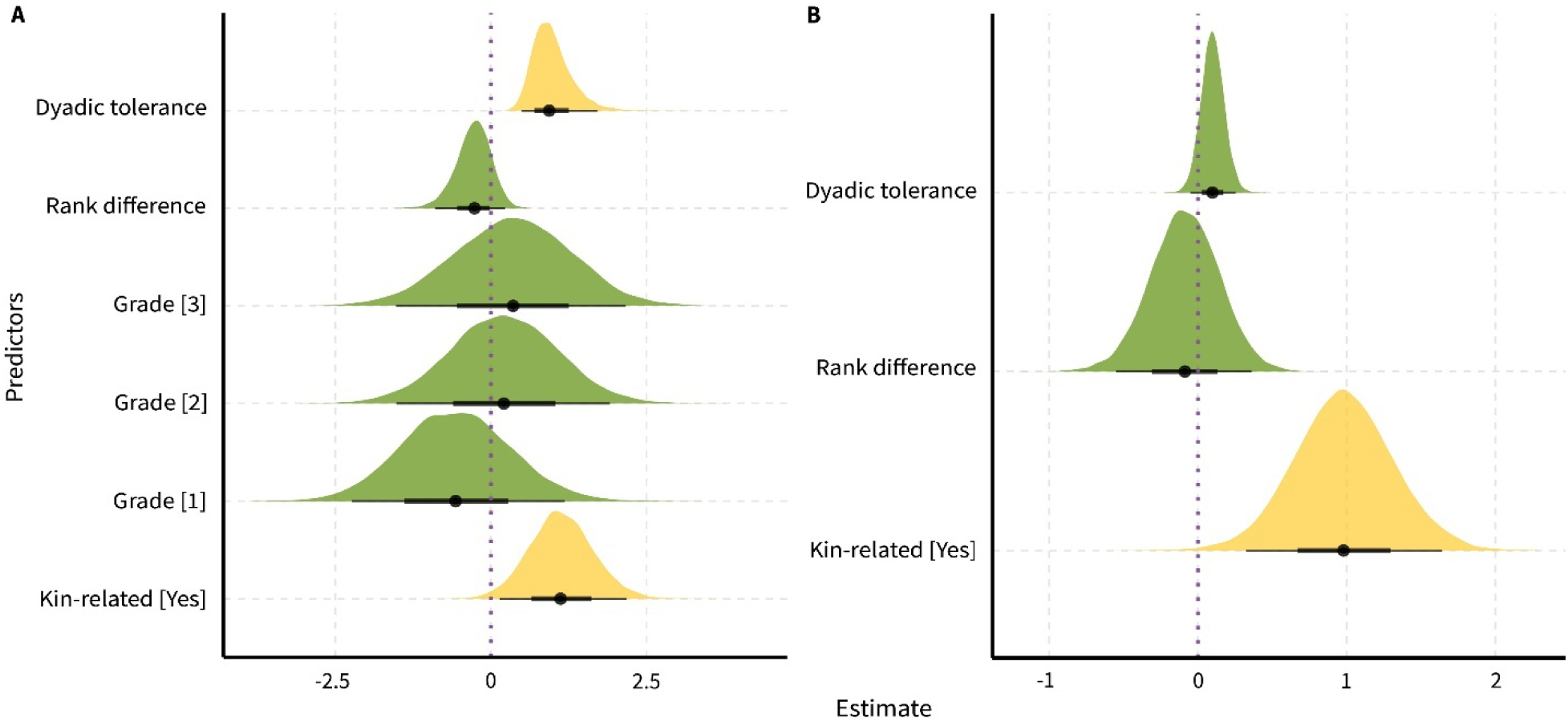
Dyadic predictors of prosocial food provisioning. **(A)** Bayesian mixed model plot showing the effects of dyadic social tolerance, rank difference, tolerance grades, and kinship on the likelihood of prosocial food provisioning. **(B)** Bayesian mixed model plot showing the effects of dyadic social tolerance, rank difference, and kinship on the magnitude of prosocial food provisioning. Yellow colors indicate robust effects of the corresponding predictors. the width of the ‘half-eye’ represents data distribution (89% credible intervals), and solid black points on the horizontal bars indicate median values. The vertical dashed lines indicate a parameter estimate of zero, i.e., the overlap of the credible interval with this line suggests no effect of the corresponding predictor.

**fig. S3.**
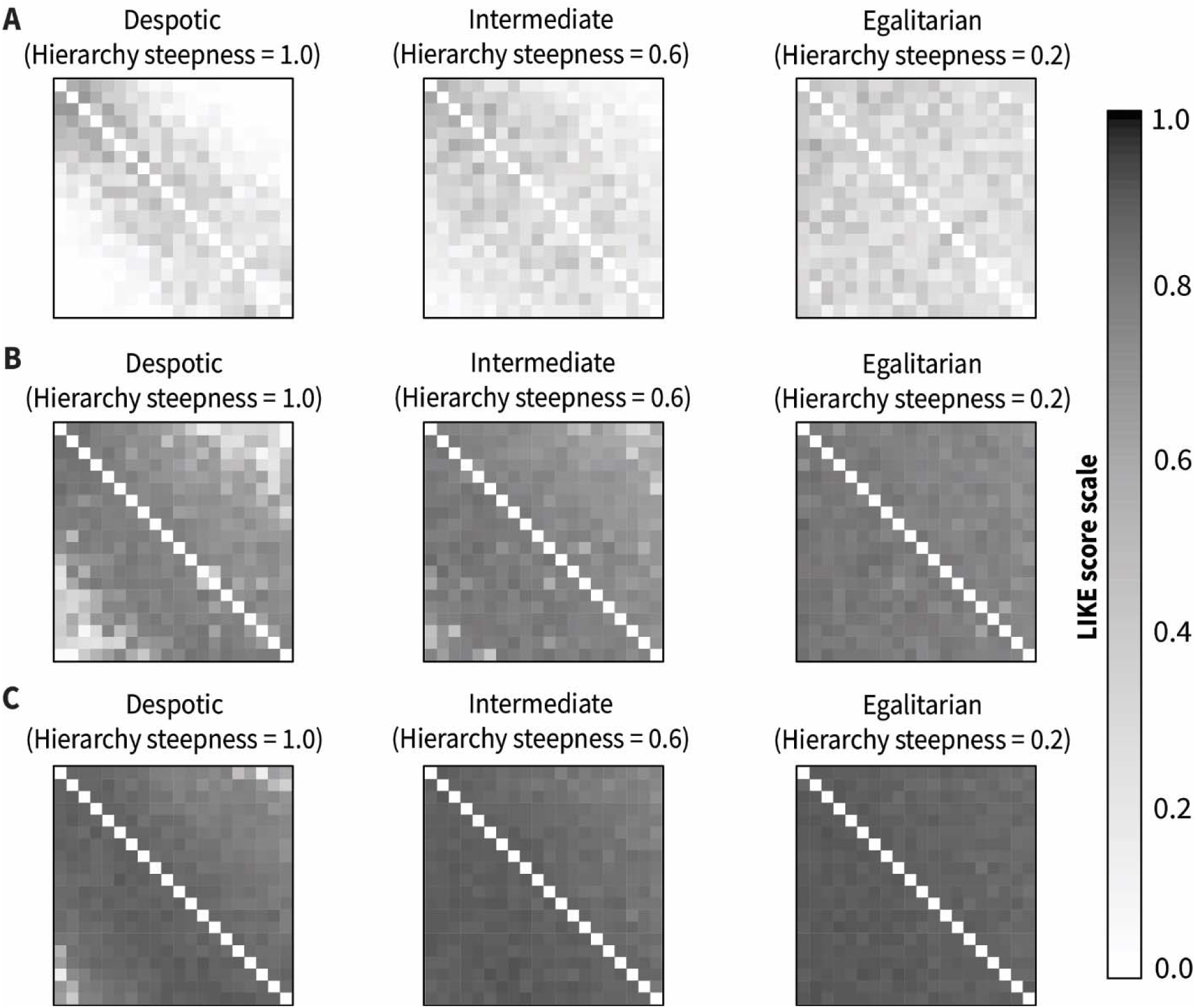
EMO-model simulation with easy-going LIKE dynamics and fast increase speed shows the emergence of LIKE relationships in societies along a despotic-egalitarian gradient. **(A)** LIKE relationships in societies with a fast decrease speed. **(B)** LIKE relationships in societies with an intermediate decrease speed. **(C)** LIKE relationships in societies with a slow decrease speed. On the y-axes, individuals are ordered from low ranking (top row) to high ranking (bottom row), and on the x-axes, from low ranking (left) to high ranking (right). Each square represents a LIKE attitude from one individual to another, indicating their LIKE relationship. A LIKE relationship ranges from 0.01 (white) to 0.99 (black), with higher values indicating stronger bonds.

**fig. S4.**
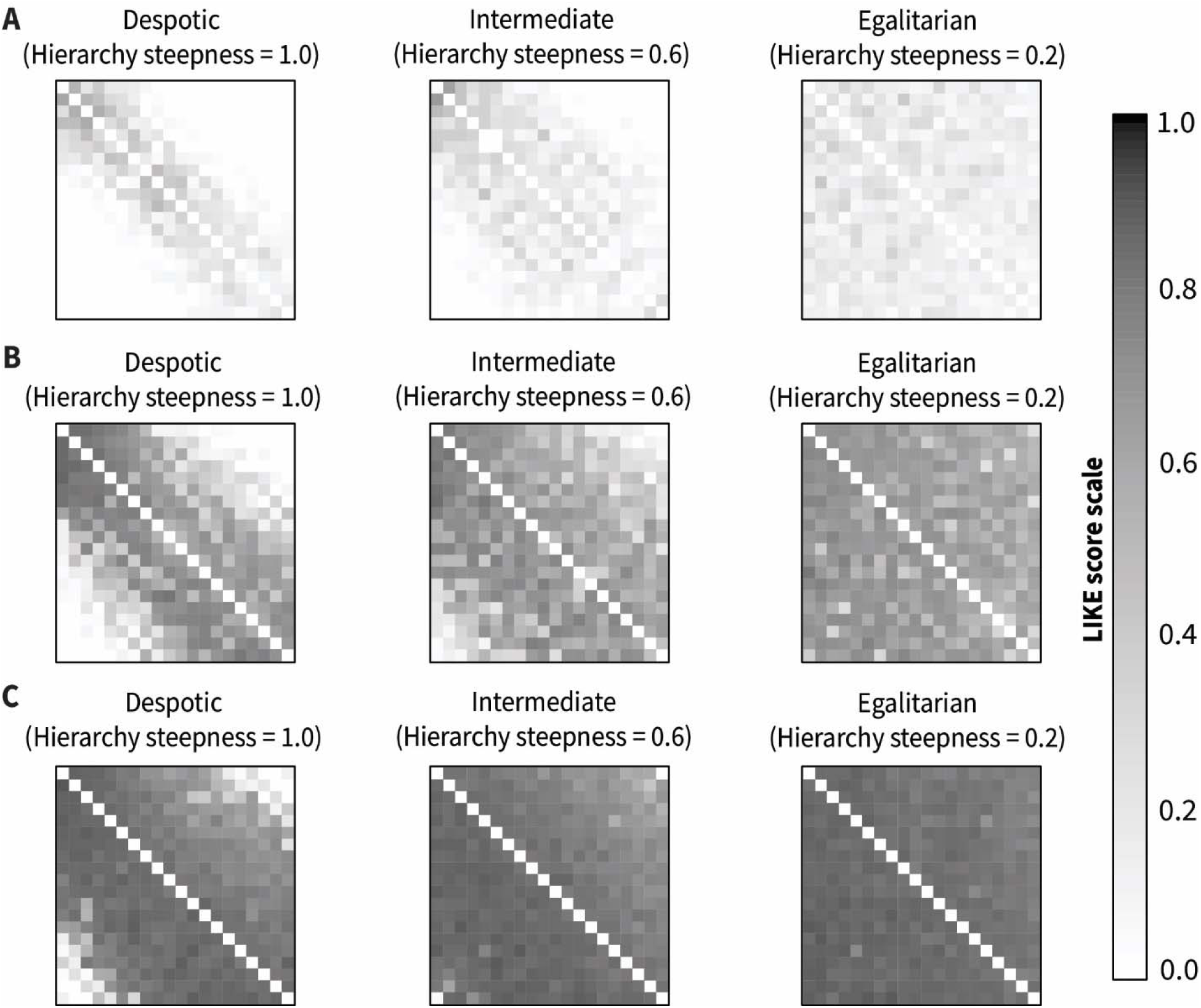
EMO-model simulation with easy-going LIKE dynamics and intermediate increase speed shows the emergence of LIKE relationships in societies along a despotic-egalitarian gradient. **(A)** LIKE relationships in societies with a fast decrease speed. **(B)** LIKE relationships in societies with an intermediate decrease speed. **(C)** LIKE relationships in societies with a slow decrease speed. On the y-axes, individuals are ordered from low ranking (top row) to high ranking (bottom row), and on the x-axes, from low ranking (left) to high ranking (right). Each square represents a LIKE attitude from one individual to another, indicating their LIKE relationship. A LIKE relationship ranges from 0.01 (white) to 0.99 (black), with higher values indicating stronger bonds.

**fig. S5.**
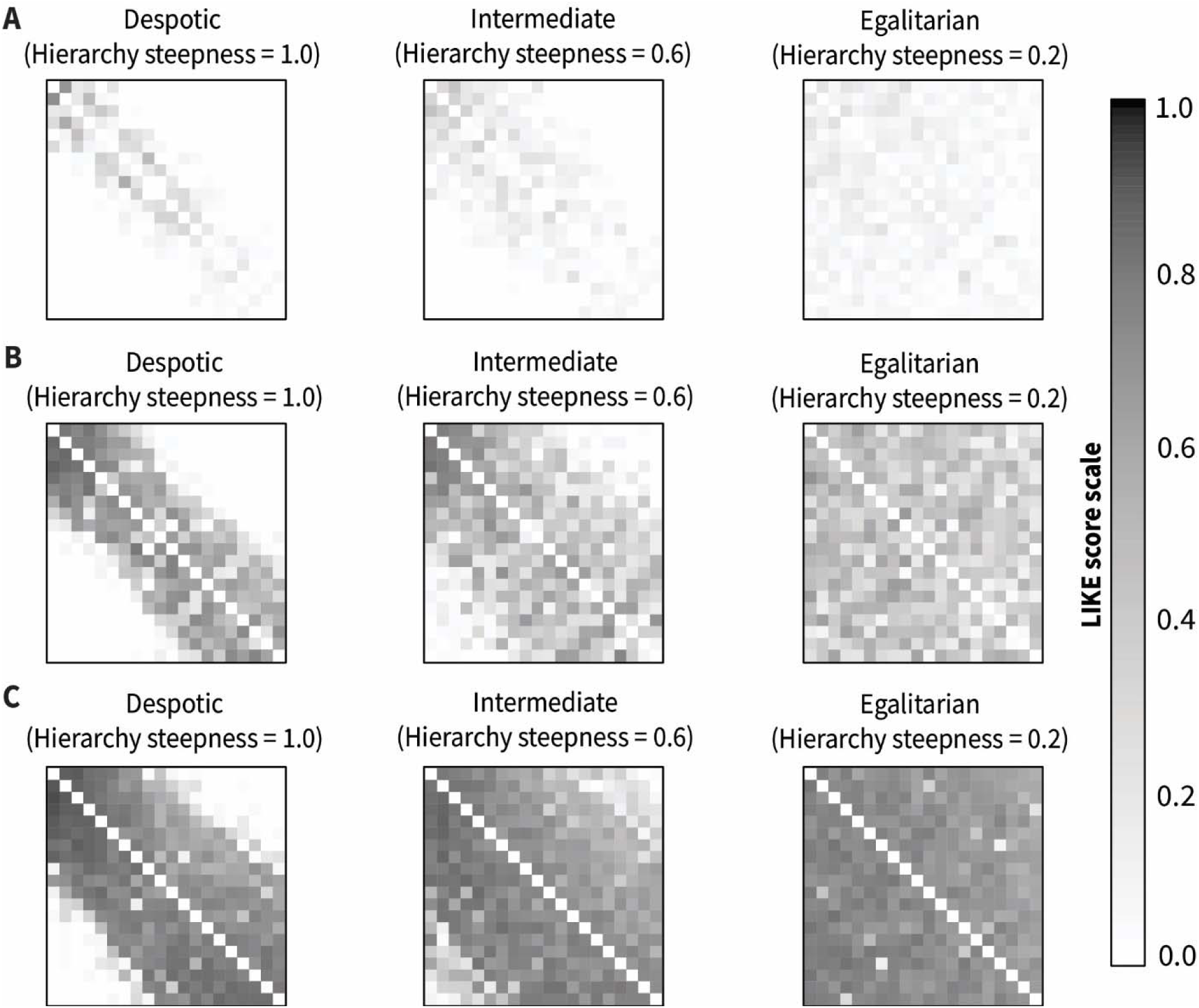
EMO-model simulation with easy-going LIKE dynamics and slow increase speed shows the emergence of LIKE relationships in societies along a despotic-egalitarian gradient. **(A)** LIKE relationships in societies with a fast decrease speed. **(B)** LIKE relationships in societies with an intermediate decrease speed. **(C)** LIKE relationships in societies with a slow decrease speed. On the y-axes, individuals are ordered from low ranking (top row) to high ranking (bottom row), and on the x-axes, from low ranking (left) to high ranking (right). Each square represents a LIKE attitude from one individual to another, indicating their LIKE relationship. A LIKE relationship ranges from 0.01 (white) to 0.99 (black), with higher values indicating stronger bonds.

**fig. S6.**
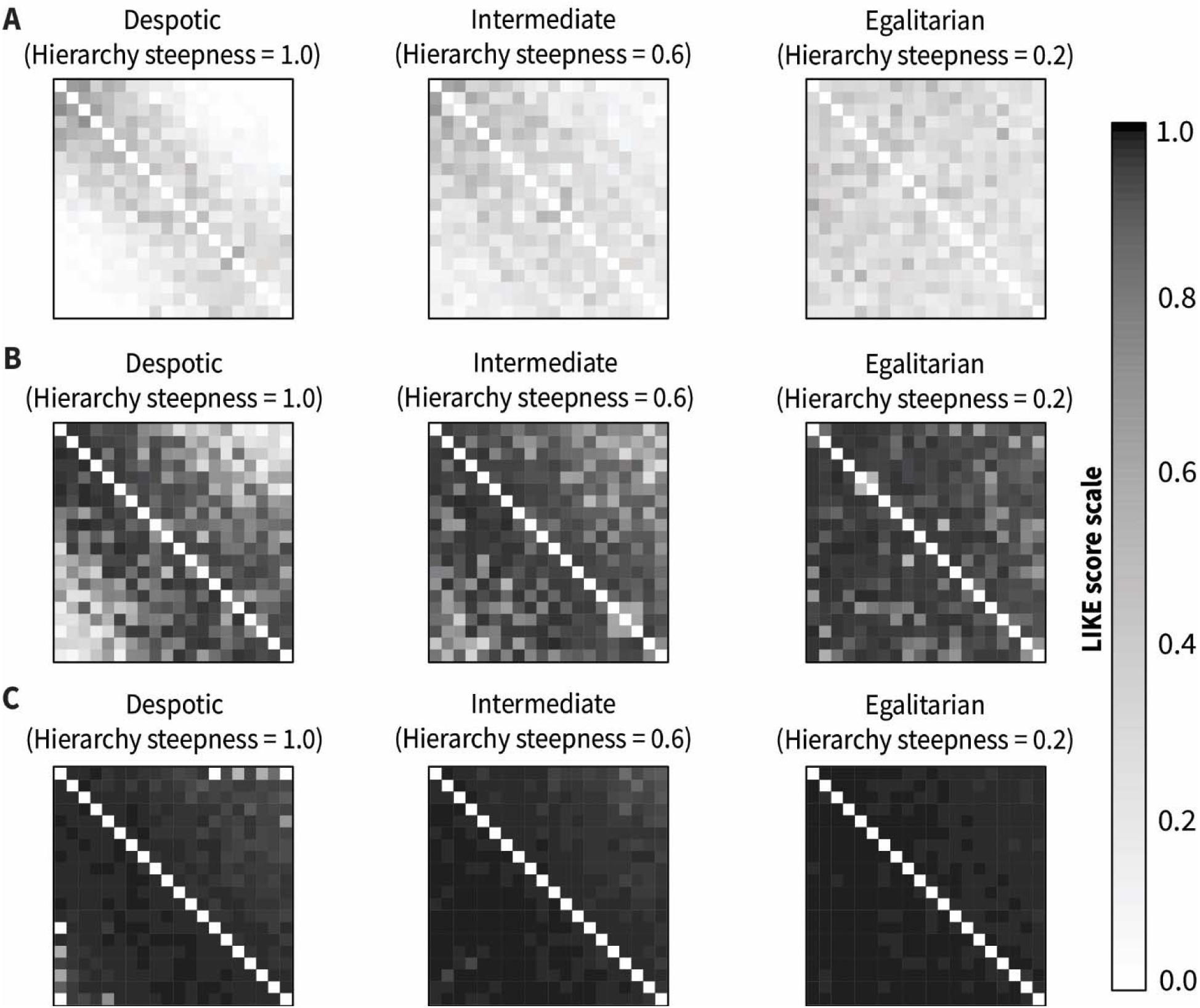
EMO-model simulation with picky LIKE dynamics and fast increase speed shows the emergence of LIKE relationships in societies along a despotic-egalitarian gradient. **(A)** LIKE relationships in societies with a fast decrease speed. **(B)** LIKE relationships in societies with an intermediate decrease speed. **(C)** LIKE relationships in societies with a slow decrease speed. On the y-axes, individuals are ordered from low ranking (top row) to high ranking (bottom row), and on the x-axes, from low ranking (left) to high ranking (right). Each square represents a LIKE attitude from one individual to another, indicating their LIKE relationship. A LIKE relationship ranges from 0.01 (white) to 0.99 (black), with higher values indicating stronger bonds.

**fig. S7.**
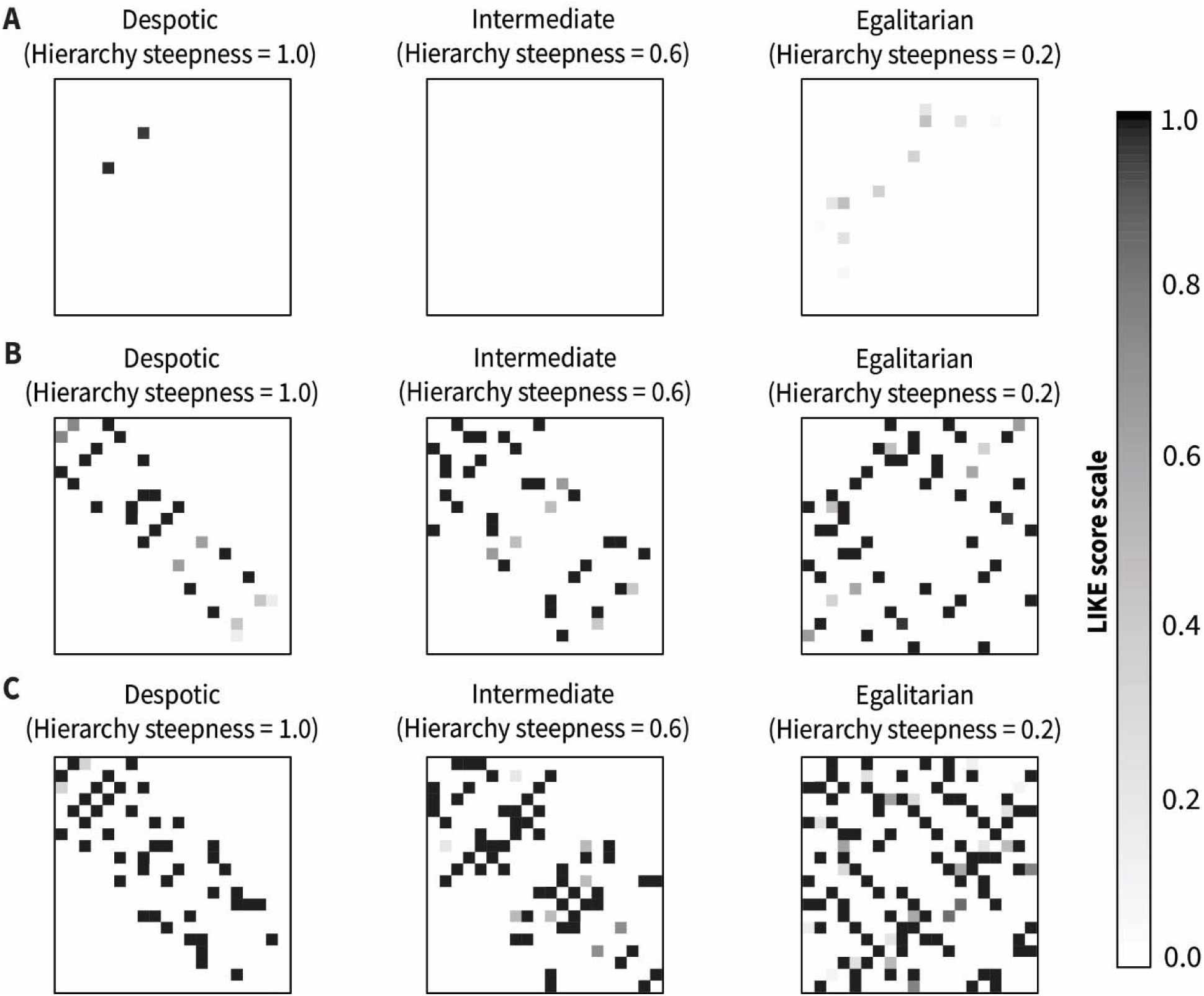
EMO-model simulation with picky LIKE dynamics and slow increase speed shows the emergence of LIKE relationships in societies along a despotic-egalitarian gradient. **(A)** LIKE relationships in societies with a fast decrease speed. **(B)** LIKE relationships in societies with an intermediate decrease speed. **(C)** LIKE relationships in societies with a slow decrease speed. On the y-axes, individuals are ordered from low ranking (top row) to high ranking (bottom row), and on the x-axes, from low ranking (left) to high ranking (right). Each square represents a LIKE attitude from one individual to another, indicating their LIKE relationship. A LIKE relationship ranges from 0.01 (white) to 0.99 (black), with higher values indicating stronger bonds.

**Table S1.** Macaque study group details and different tests performed.

| Macaque Species | Group Name | Observations and Tests |  |  |  |
| --- | --- | --- | --- | --- | --- |
|  |  | Observations | Cooperation | Prosociality | Co-feeding |
| *Macaca fuscata* (Japanese) | Affenberg |  | F=13, M=3, J=9 ,<br>A=7 | F=19, M=6, J=9,<br>A=16; # (13) |  |
| *Macaca mulatta* (Rhesus) | R3G2 | N=15; 1300 min<br>(86.66±14.47) min | F=6, M=1, J=3,<br>A=4 |  | N=15, F=13,<br>M=2, J=3, A=12 |
|  | R3G7 | N=14, 2947.4 min<br>(210.53±18.13) min | F=2, J=2 | F=13, M=1, J=4,<br>A=10 | N=13, F=12,<br>M=1, J=3, A=10 |
| *Macaca fascicularis* (Long-tailed) | J1G4 | N=11; 3520 min<br>(320) min | F=5, M=2, J=2,<br>A=5; # (12) | F=9, M=2, J=4,<br>A=7; # (27) | N=11, F=9,<br>M=2, J=3, A=8 |
|  | J1G7 | N=17; 5440 min<br>(320) min | F=6, M=4, J=5,<br>A=5; # (12) | F=13, M=2, J=6,<br>A=9; # (27) | N=17, F=12,<br>M=5, J=7, A=10 |
|  | NWR | N=4; 1280 min<br>(320) min | M=4, A=4; # (12) | M=4, A=4; # (27) | N=4, M=4, A=4 |
| *Macaca silenus* (Lion-tailed) | Blijdorp | N=3; 740 min<br>(246.66±30.55) min | F=1, M=1, A=2 | F=3, M=1, A=4 | N=3, F=2, M=1,<br>A=3 |
|  | Apenheul | N=8; 2820 min<br>(352.5±10.35) min |  |  | N=8, F=5, M=3,<br>J=2, A=6 |
| *Macaca sylvanus* (Barbary) | Gaia | N=14; 3410.7 min<br>(243.62±1.65) min | F=1, M=4, A=5 | F=8, M=6, A=14 | N=14, F=8,<br>M=6, A=14 |
|  | Apenheul | N=9; 3855 min<br>(428.33±7.81) min |  |  | N=9, F=8, M=1,<br>A=9 |
| *Macaca nigra* (Crested) | Blijdorp | N=5; 2353.7 min<br>(470.75±114.44) min | F=1, M=3, J=2,<br>A=2 | M=3, F=2, J=2,<br>A=3 | N=6, F=3, M=3,<br>J=3, A=3 |
|  | Artis | N=4; 1280 min<br>(320) min | F=2, M=1, J=1,<br>A=2 | F=3, M=1, J=1,<br>A=3 |  |
|  | Planckendael | N=5; 909 min<br>(181.88±25.84) min |  |  | N=5, F=2, M=3,<br>A=5 |

*Note:* Colored cells indicate observations or tests performed, and blank cells indicate that observations or tests could not be performed due to COVID-19 restrictions. N = Sample size, F = Female, M = Male, J = Juvenile and sub-adults (>1 year but ≤ 3.5 years at the time of testing), A = Adults (>3.5 years). In the observations column, total observation minutes are given along with information on mean ± standard deviation per individual in parentheses. For cooperation, the numbers indicate the self-trained participating individuals (cf. **Data S1**). References (#) show the use of published datasets for these tests using experimental designs identical to the current study.

**Table S2.** Effects of dyadic tolerance, rank difference, tolerance grades, prosociality, and kinship on the likelihood of cooperation.

|  | Estimate | 89% critical interval | Probability of direction |
| --- | --- | --- | --- |
| Dyadic tolerance | 0.86 | 0.40, 1.37 | 0.99 |
| Rank difference | -0.37 | -0.83, 0.07 | 0.91 |
| Grade [1] | -0.28 | -1.64, 1.10 | 0.63 |
| Grade [2] | -0.12 | -1.45, 1.23 | 0.55 |
| Grade [3] | -0.28 | -1.69, 1.14 | 0.62 |
| Prosocial [Yes] | 1.41 | 0.58, 2.27 | 0.99 |
| Kinship [Yes] | 1.23 | 0.29, 2.18 | 0.98 |

*Note:* only best-fitted model is reported.

**Table S3.** Effects of dyadic tolerance, rank difference, tolerance grades, prosociality, and kinship on the magnitude of cooperation.

|  | Estimate | 89% critical interval | Probability of direction |
| --- | --- | --- | --- |
| Dyadic tolerance | 0.23 | 0.09, 0.40 | 0.99 |
| Rank difference | -0.17 | -0.46, 0.11 | 0.83 |
| Grade [1] | -0.51 | -1.68, 0.72 | 0.76 |
| Grade [2] | 0.24 | -0.96, 1.35 | 0.64 |
| Grade [3] | 0.37 | -0.85, 1.56 | 0.69 |
| Prosocial [Yes] | 0.46 | -0.11, 1.03 | 0.9 |
| Kinship [Yes] | -0.45 | -1.03, 0.13 | 0.89 |

*Note:* only best-fitted model is reported.

**Table S4.** Effects of tolerance grades, group size, groom index, and aggression index on the likelihood of cooperation.

|  | Estimate | 89% critical interval | Probability of direction |
| --- | --- | --- | --- |
| Grade [1] | -0.20 | -1.64, 1.23 | 0.59 |
| Grade [2] | 0.11 | -1.21, 1.44 | 0.55 |
| Grade [3] | -0.24 | -1.60, 1.12 | 0.61 |
| Group size | -0.26 | -0.48, -0.07 | 0.98 |
| Groom index | 0.17 | -0.30, 0.63 | 0.73 |
| Aggression index | 0.47 | -0.00, 0.96 | 0.94 |

*Note: only best-fitted model is reported.*

**Table S5.** Effects of tolerance grades, group size, groom index, and aggression index on the magnitude of cooperation.

|  | Estimate | 89% critical interval | Probability of direction |
| --- | --- | --- | --- |
| Grade [1] | 0.27 | -1.07, 1.60 | 0.63 |
| Grade [2] | 0.27 | -0.98, 1.44 | 0.65 |
| Grade [3] | 0.18 | -1.08, 1.41 | 0.59 |
| Group size | -0.07 | -0.19, 0.04 | 0.84 |
| Groom index | -0.11 | -0.42, 0.23 | 0.70 |
| Aggression index | 0.26 | -0.07, 0.60 | 0.90 |

*Note: only best-fitted model is reported.*

**Table S6.** Effects of dyadic tolerance, rank difference, tolerance grades, and kinship on the likelihood of prosocial food provisioning.

|  | Estimate | 89% critical interval | Probability of direction |
| --- | --- | --- | --- |
| Dyadic tolerance | 0.98 | 0.57, 1.53 | 1.00 |
| Rank difference | -0.28 | -0.76, 0.14 | 0.85 |
| Grade [1] | -0.56 | -1.93, 0.86 | 0.74 |
| Grade [2] | 0.21 | -1.18, 1.59 | 0.59 |
| Grade [3] | 0.35 | -1.16, 1.82 | 0.65 |
| Kinship [Yes] | 1.13 | 0.33, 19.6 | 0.98 |

*Note: only best-fitted model is reported.*

**Table S7.** Effects of dyadic tolerance, rank difference, and kinship on the magnitude of prosocial food provisioning.

|  | Estimate | 89% critical interval | Probability of direction |
| --- | --- | --- | --- |
| Dyadic tolerance | 0.10 | -0.02, 0.22 | 0.90 |
| Rank difference | -0.09 | -0.46, 0.28 | 0.64 |
| Kinship [Yes] | 0.98 | 0.45, 1.51 | 0.99 |

*Note: only best-fitted model is reported.*

## Acknowledgments

We thank all the staff members of the Affenberg Zoobetriebsgesellschaft mbH (Austria), the Biomedical Primate Research Centre (the Netherlands), Diergaarde Blijdorp (the Netherlands), Gaia Zoo (the Netherlands), Apenheul Primate Park (the Netherlands), Planckendael Zoo (Belgium), and Artis Zoo (the Netherlands) for their support and assistance during the research work. We sincerely thank Peter Gaubatz, Svenja Gaubatz, Max Dorner, Jan A.M. Langermans, Annet Louise Louwerse, Jos Hartog, Linda Bruins-van Sonsbeek, Emile F. Prins, Lisette van den Berg, and Marjolein Osieck for allowing us to conduct the studies at the different zoos and institutions. We thank Sjoerd Sijbrandij, Paola Meems, Amber Kozanli, Yesper Bos, and Mary Maximiadi for their help with data organization. We thank Sudeshna Chakraborty, Julia Ostner, Oliver Schülke, and Thomas Bugnyar for their feedback on earlier versions of the manuscript.

## Author contributions

DB and JJMM conceptualized the study; DB, TWZ, EJCvL, and JJMM developed the methodology; DB, TSR, and TWZ analyzed the data with input from EJCvL and JJMM; DB, TSR, TWZ, and VS designed and edited the figures with input from JJMM; DB, EB, SC, PEC, EC, JAdJ, EJAM, ARG, EJJ, CEK, PENK, EM, VIS, ESJvD, JV, SW, and AZ collected and coded the data. DB and JJMM secured funding for this work; DB wrote the original manuscript with input from JJMM. DB, TWZ, TSR, KRLJ, LSP, EHMS, EJCvL, and JJMM edited the manuscript. All authors read and approved the manuscript.

## Data and materials availability

Data and code will be made publicly available upon publication.

